# Quantification of translation uncovers the functions of the alternative transcriptome

**DOI:** 10.1101/608794

**Authors:** Lorenzo Calviello, Antje Hirsekorn, Uwe Ohler

**Affiliations:** Berlin Institute for Medical Systems Biology, Max-Delbrück-Center for Molecular Medicine, Hannoversche Strasse 28, Berlin 10115, Germany; Department of Biology, Humboldt-Universität zu Berlin, Unter den Linden 6, Berlin 10117, Germany; Department of Computer Science, Humboldt-Universität zu Berlin, Unter den Linden 6, Berlin 10117, Germany; University of California San Francisco, Department of Cell & Tissue Biology, 513 Parnassus Avenue, 94143, San Francisco, California, United States

## Abstract

At the center of the gene expression cascade, translation is fundamental in defining the fate of much of the transcribed genome. RNA sequencing enables the quantification of complex transcript mixtures, often detecting several splice isoforms of unknown functions for one gene. We have developed *ORFquant*, a new approach to annotate and quantify translation at the single open reading frame (ORF) level, using information from Ribo-seq data. Relying on a novel approach for transcript filtering, we quantify translation on thousands of ORFs, showing the power of Ribo-seq in revealing alternative ORFs on multiple isoforms per gene. While we find that one ORF represents the dominant translation product for most genes, we also detect genes with translated ORFs on multiple transcript isoforms, including targets of RNA surveillance mechanisms. Assessing the translation output across human cell lines reveals the extent of gene-specific differences in protein production, which are supported by steady-state protein abundance estimates. Computational analysis of Ribo-seq data with *ORFquant* (available at https://github.com/lcalviell/ORFquant) provides a window into the heterogeneous functions of complex transcriptomes.

## Introduction

Studying gene expression allows us to understand the functions of different molecules and regulatory sequence elements, whether they act at the level of transcription, the transcribed RNA, or the encoded protein. To ensure correct protein synthesis, transcriptional and post-transcriptional regulatory programs determine the identity and amount of mature RNA templates. The translation process ensures the correct identity and amount of synthesized proteins.

The ribosome is the main actor of the translation process, a complex ribonucleoparticle that is not only able to synthesize proteins, but also acts as quality control platform for both the nascent peptide^1^ and the translated mRNA^2^. Several RNA surveillance mechanisms are known to occur co-translationally, and their importance for different processes such as differentiation or disease has been investigated^3^.

Ribosome profiling (Ribo-seq) has made it possible to pinpoint the positions of actively translating ribosomes transcriptome-wide, using ribosome footprinting coupled to RNA sequencing^4^. In the last decade, Ribo-seq has been extensively used to investigate the molecular mechanisms acting on the ribosome, and to identify the entire ensemble of translated regions (the *translatome*) in multiple organisms and conditions. The resulting rich datasets have triggered a plethora of dedicated analysis methods, which exploit distinct features of Ribo-seq profiles to confidently identify translated ORFs^5,6^. In this context, many reports have focused on whether small translated regions are hidden in long non-coding RNAs^7–9^, with less attention given so far to account for the presence of multiple transcript isoforms per gene.

Transcript diversity can result from either alternative splicing (AS) or from alternative transcription start or poly-adenylation site usage, and it is now commonly profiled by RNA-seq experiments, which measure steady-state abundance of (m)RNAs. Large-scale efforts have uncovered the wide spectrum of alternative transcript isoforms, with many being lowly expressed and/or presenting incomplete ORFs^10^. The contribution of this transcript heterogeneity to an expanded translatome is therefore an intensely debated topic^11,12^, with much of transcript and protein abundance apparently explained by a single dominant transcript per gene^13^.

The mere presence of multiple transcripts does not imply the presence of a distinct, functional protein translated from each transcript isoform: transcripts might be retained in the nucleus, selectively degraded, or undergo translational repression. From a technical point of view, RNA-seq experiments quantify a complex scenario in which, depending on the protocol used, alternative transcripts may also reflect different steps of RNA processing and not the stable, steady-state cytoplasmic pool of mRNAs available to the ribosome. From a different direction, shotgun proteomics approaches are only recently providing the sensitivity to detect tens of thousands of proteins from a single sample^14^ and rarely reach the depth to investigate alternative protein isoforms.

To close this gap, we developed a strategy to identify and quantify translation on the subset of transcripts that are expressed in the cell. A recent study presented a proof-of-principle for validating the presence of multiple transcript isoforms in Ribo-seq data^15^, underlining the potential of isoform-aware analysis approaches to fully define the translatome. Following up on this premise, we here describe *ORFquant*, a Ribo-seq analysis approach that detects and quantifies ORF translation across multiple transcript isoforms and zooms in on the potential roles of alternative transcripts.

## Results

### The ORFquant approach to annotate and quantify translation

Our approach is based on the premise that, despite their short length, Ribo-Seq reads are sufficient to support a given set of alternative transcripts (Figure 1a, b). Single-nucleotide positions corresponding to the peptidyl-site for each ribosome (P-sites positions) and junction reads are first extracted from the Ribo-seq alignment (Methods) and then mapped to flattened gene models from a given annotation (Figure 1b). In this way, transcript features (e.g. exonic bins or splice junctions) are designated as unique or shared across multiple annotated transcript isoforms.

**Figure 1:**
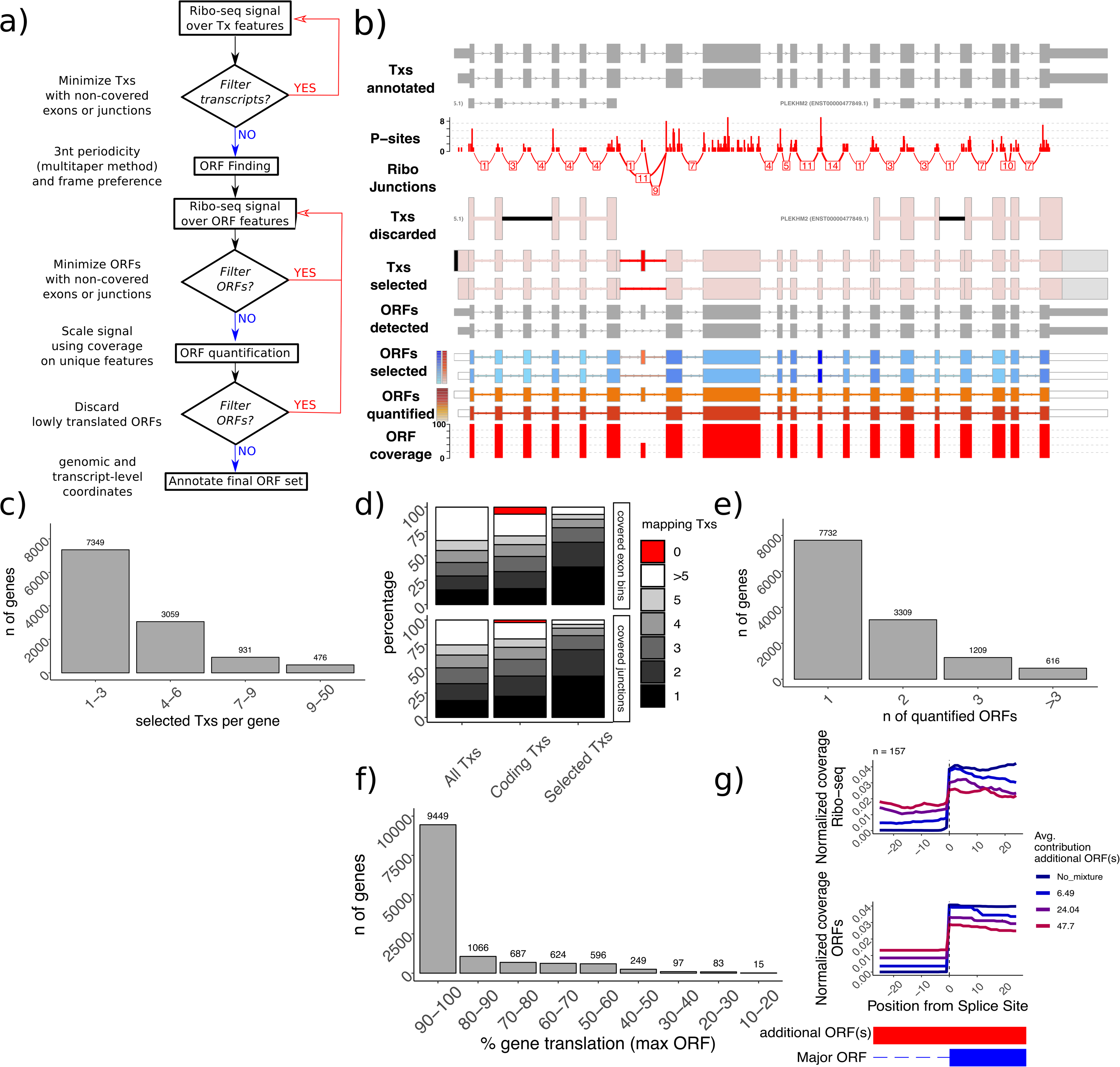
The *ORFquant* strategy to quantify translation on selected transcripts. a) The *ORFquant* workflow; b) the *PHLEKM2* gene as an example: displayed tracks represent, from top to bottom: 1) complete annotation, 2) P-sites positions, 3) junction reads (from Ribo-seq), 4) discarded transcripts, 5) selected transcripts, 6) detected ORFs, 7) selected ORFs, 8) quantified ORFs and 9) ORF coverage (defined here using the percentage of gene translation). Colors for discarded and selected transcripts indicate unique features with no signal (*black*); shared features with no signal (*grey*); unique features with signal (*red*); and shared features with signal (*pink*). Colors for discarded and selected ORFs indicate signal in shared features (*blue heatmap*) and signal in unique features (*red heatmap*). For the quantified ORFs, the heatmap indicates ORF coverage values (0-100). Thick bars indicate CDS regions, as defined by the annotation or by *ORFquant* (de-novo). c) Number of selected transcripts per gene (*x-axis*) against number of genes (*y-axis*). d) Percentage of covered junctions (*bottom*), or covered exons (*top*) mapping to a different number of structures using all transcripts, protein-coding transcripts only or selected transcripts only. e) The number of quantified ORFs (*x-axis*) is shown against number of genes (y-axis). f) The number of genes (*y-axis*) are plotted against the contribution (in percentages) of their major ORF. g) Aggregate plot of Ribo-seq coverage (normalized 0-1 per each region) and ORF coverage (*ORF_pct_P_sites_pN*, Methods) over candidate alternative splice sites regions as detected by *ORFquant*. No mixture indicates one ORF only, while other tracks indicate the presence of additional ORFs, divided by their summed translation values. Explanatory scheme at the bottom, with blue representing the major ORF and red the additional ORF(s).

We first retain a subset of annotated transcripts, which is sufficient to explain all the observed P-sites or junction reads and reduce the occurrence of exons and junctions with no signal, using an Occam’s razor strategy (Methods). In brief, a transcript is filtered out if its features supported Ribo-seq signal can be explained by another transcript with better support (i.e. containing more features with Ribo-seq support or fewer unsupported features). As Ribo-seq reads are largely found in 5’UTRs and coding regions only, this approach might not distinguish between transcripts differing in their 3’UTR.

This simple yet effective selection strategy leads to a significantly reduced number of transcripts: the observed Ribo-seq signal can be explained by 1 to 3 transcript structures for most genes, without showing a strong bias for expression level (Figure 1c, Supplementary Figure 1). This selection dramatically improves the assignments of both exons and junctions to transcripts (Figure 1d): when considering covered exons or junctions (defined as having at least one Ribo-seq read mapped to them), ~64% of exons mapped to 1 or 2 transcripts, compared to ~29% when no selection is performed. Considering only annotated protein-coding transcripts does not substantially improve the mapping of covered features, while it ignores the presence of covered exons and junctions unique to non-coding transcripts. Next, we detect translated ORFs *de novo* in each of the selected transcripts, using frame preference and the multitaper^16–18^ test to select in-frame signal displaying 3nt periodicity (Methods), a hallmark of active translation elongation. Detected ORFs are filtered using the same strategy used for transcript filtering.

After calculating coverage on unique and shared ORF features (exonic bins and splice junctions within ORF boundaries), a scaling factor between 0 and 1 is determined using the coverage on unique ORF features, or the amount of overlap between ORFs when no unique feature can be detected (Methods). This scaling factor represents the fraction of Ribo-seq signal which can be assigned to that ORF. The scaled number of P-sites is then normalized by the ORF length to arrive at transcripts per million (TPM)^19^-like values, named ORFs per Million (“*ORFs_pM”*). Moreover, we calculate the relative contribution of each ORF to the overall translation output of each gene (“*ORF_pct_P_sites”*, or percentage of gene translation). An additional filtering step discards poorly translated ORFs. ORFs are then annotated according to their position relative to their host transcript, to other detected ORFs in the same gene, and to annotated CDS regions.

Applying *ORFquant*, we quantified translation for ~20,800 ORFs in ~12,300 genes profiled in a Ribo-Seq data set from the human HEK293 cell line^18^. Most genes (7,732 Figure 1e) displayed only one translated ORF, with another >5,000 genes showing translation of multiple ORFs. Upon closer inspection (Figure 1f), we observed that for the majority of genes (~80%), the most translated (i.e., major) ORF could explain >80% of the total gene translation, with only 444 genes for which the major ORF explained <50% of the translational output. We did not observe a clear dependency between number of detected ORFs (or % of translation of major ORF) and overall Ribo-seq coverage, with the exception of the few dozen genes for which the major ORF accounted for little of the total gene output (Supplementary Figure 1).

In principle, the final set of ORFs can be provided to any algorithm for transcript quantification. To demonstrate the effectiveness of our simple approach, we compared our estimates with the ones calculated by RSEM^19^, a well-known statistical approach devoted to transcript quantification. We observed good correlation between the two method in their estimates of the relative contribution of each ORF to the total output (Supplementary Figure 2, *left*). In addition, we observed how RSEM quantification estimates showed high uncertainty (Methods) for ORFs where few unique features are present, which are cases where *ORFquant* assigns low translation estimates to the major ORF (Supplementary Figure 2, *right*). Despite major differences in their quantification strategy (Discussion), both *ORFquant* and RSEM showed similar performances in determining the contribution of each ORFs to the total gene translation output. To illustrate the consistency of our translation estimates, we annotated the ORF structures with respect to the major (most translated) ORFs in each gene: this allowed us to detect genomic regions (e.g. different alternative splice sites) where the Ribo-seq signal should reflect different quantitative estimates of translation coming from different ORF(s). Aggregate profiles of Ribo-seq coverage closely reflected the expected pattern calculated by *ORFquant* (Figure 1g). Additional profiles over different genomic locations are shown in Supplementary Figure 3. Taken together, the translation of a major ORF accounts for >80% of total gene translation for most of the genes, but distinct translated ORFs are detected from multiple translated transcripts for hundreds of genes.

### Quantification of translation as a window into the functional relevance of alternative open reading frames

As translation is a cytoplasmic process, we expected the ensemble of transcript structures selected by *ORFquant* to represent *bona fide* cytoplasmic transcripts. To test this hypothesis, we performed a differential exon usage analysis^20^, using RNA-seq data from nuclear and cytoplasmic extracts in HEK293 cells^21^. Most exons unique to discarded structures showed marked nuclear localization (log2FC>0), while exons of selected transcripts showed a prominent cytoplasmic enrichment (Figure 2a). Translated transcripts displayed a more marked cytoplasmic localization. An example of the selection strategy discarding pre-mRNA structures in favor of cytoplasmic transcripts is shown in Figure 2b.

**Figure 2:**
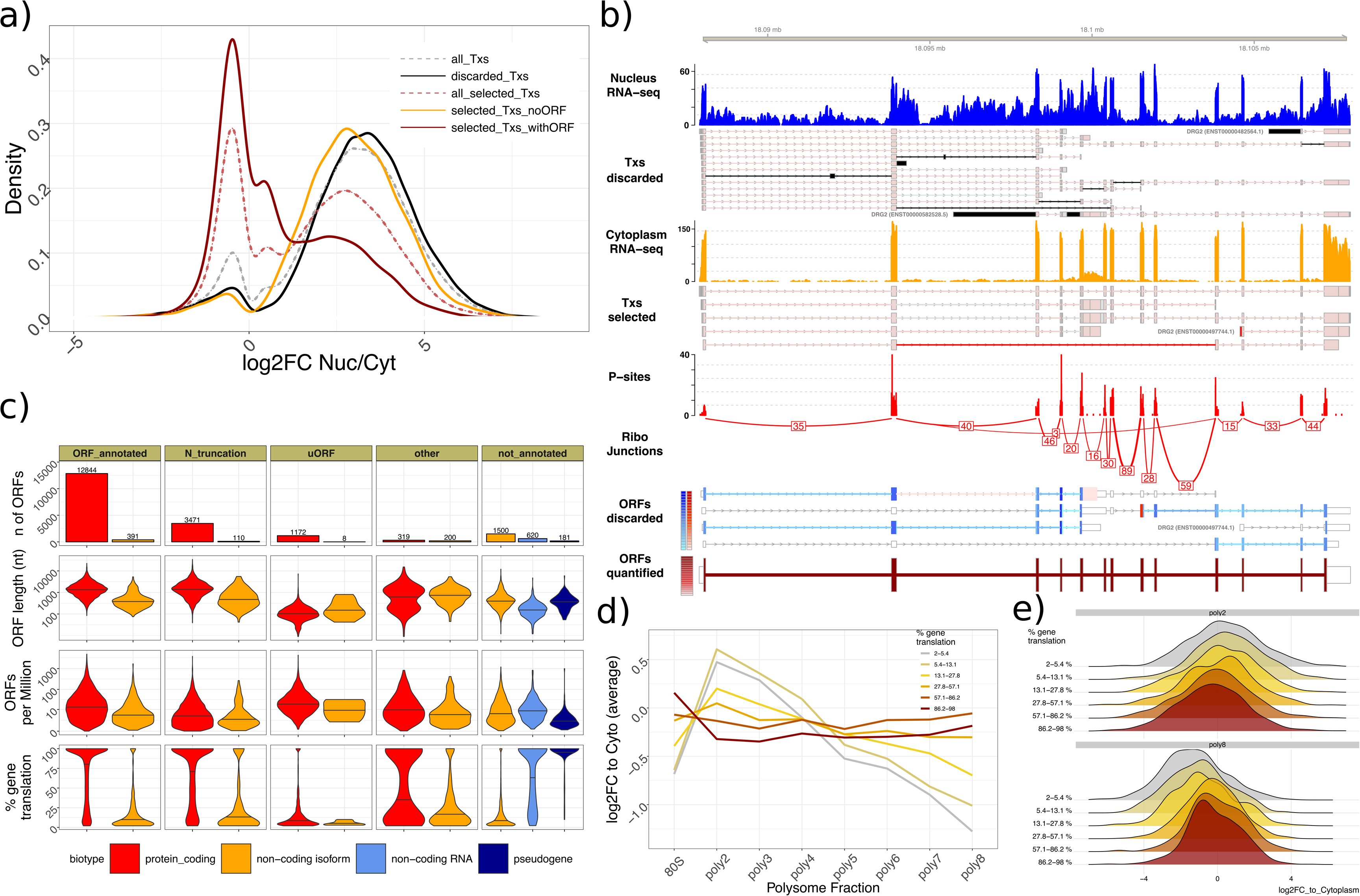
Quantification of translation on cytoplasmic mRNAs. a) Density of exonic fold changes for nuclear and cytoplasmic RNA-seq for different transcript classes. Negative values indicate more cytoplasmic abundance, while positive values indicate enrichment in the nucleus. b) The *DGR2* locus as example: tracks represent, in descending order: 1) Nuclear RNA-seq coverage, 2) discarded transcripts, 3) cytoplasmic RNA-seq coverage, 4) selected transcripts, 5) P-sites positions, 6) junction reads, 7) discarded ORFs, 8) quantified ORFs. Color representation as in Figure 1b. c) Overview of the *ORFquant*-derived translatome. Number of ORFs, ORF length in nucleotides, length-normalized quantification and % of gene translation are shown, stratified by ORF category and annotated biotype. *ORF_annotated* represents ORFs whose structure perfectly matches the annotated CDS; *other* represents additional ORFs, such as nested ORFs, overlapping ORFs, downstream ORFs in 3’UTRs (dORFs), while *not_annotated* represents ORFs in transcripts with no CDS annotation. The maximum width of each violin plot is the same for each panel, and the median value is shown as a black bar. d) Average exonic fold changes with respect to cytoplasmic abundance (y-axis) for different polysome fractions (x-axis) for ORFs exhibiting different levels of translation within the same genes. e) Density plot of aforementioned exonic fold changes for two polysome fractions and for different ORF classes.

When examining the GENCODE annotation^22^ of the transcripts hosting *de novo* identified ORFs, we noticed ~2,000 ORFs in non-coding transcript isoforms of protein-coding genes, most of which lacked annotated ORFs (Figure 2c). Compared to ORFs in annotated protein-coding transcript isoforms, these ORFs exhibited much lower translation, accounting for a median of 6.8 % of gene translation, compared to 87% for ORFs that fully matched annotated CDSs. More than 3,500 N-terminal truncation events were also detected, showing high levels of translation. Upstream ORFs (uORFs) and other small ORFs exhibited low signal, albeit high when normalized by their length. In annotated non-coding genes, we detected 181 ORFs from annotated pseudogenes and 620 ORFs from other non-coding RNA genes, with overall lower translation levels than protein-coding RNAs (Figure 2c).

Analysis of a deep polysome profiling dataset (Trip-Seq^23^) from the same cell line showed that the quantitative estimates of translation agreed with distinct polysome profiles (Figure 2d,e; Supplementary Figure 4): Exons uniquely mapping to transcripts harboring lowly translated ORFs accumulated in low polysomes and were depleted in heavier polysomal fractions. Conversely, highly translated transcripts exhibited sustained levels also in heavy polysomes. Despite the fundamental differences between polysome profiling and Ribo-seq in representing the translated transcriptome, the two techniques therefore agreed in detecting quantitative differences in the translation of multiple transcripts per gene.

The presence of numerous lowly translated ORFs detected in non-coding transcript isoforms (Figure 2c) suggested inefficient translation and/or low steady-state abundance of the translated transcript. We wondered whether transcripts subject to RNA surveillance mechanisms (such as nonsense-mediated decay, NMD) might cause such a low but detectable Ribo-seq signal. The presence of a premature termination codon (PTC) is an important feature of many NMD targets^24^, which is assumed to be recognized as such when the stop codon is located sufficiently upstream of the last splice junction, i.e. when a downstream Exon Junction Complex (EJC) is not displaced during translation elongation (Figure 3a). To investigate the putative action of NMD on PTC-containing transcripts, we divided transcripts based on the presence of a splice site downstream of a detected ORF. A recent study mapped NMD-mediated cleavage events on the transcriptome in HEK293 cells^25^, by knocking down *XRN1*, the exonuclease in charge of degrading the cleaved transcripts. When aligning the cleavage sites at the stop codons of (putative) PTC- and non-PTC-containing transcripts (from the same genes), we observed a clear difference (Figure 3b): transcripts without PTC, i.e. where all EJCs are presumably displaced, showed background-like signal, while transcripts harboring a putative PTC showed a marked degradation profile around their stop codon^24^. The degradation signal was less pronounced when *SMG6* or *UPF1* were also knocked-down, underlining the effect of known key factors of the NMD pathway on our candidate NMD targets. A clear example of such pattern is visible on a translated ORF in the *SNHG17* gene (Figure 3c).

**Figure 3:**
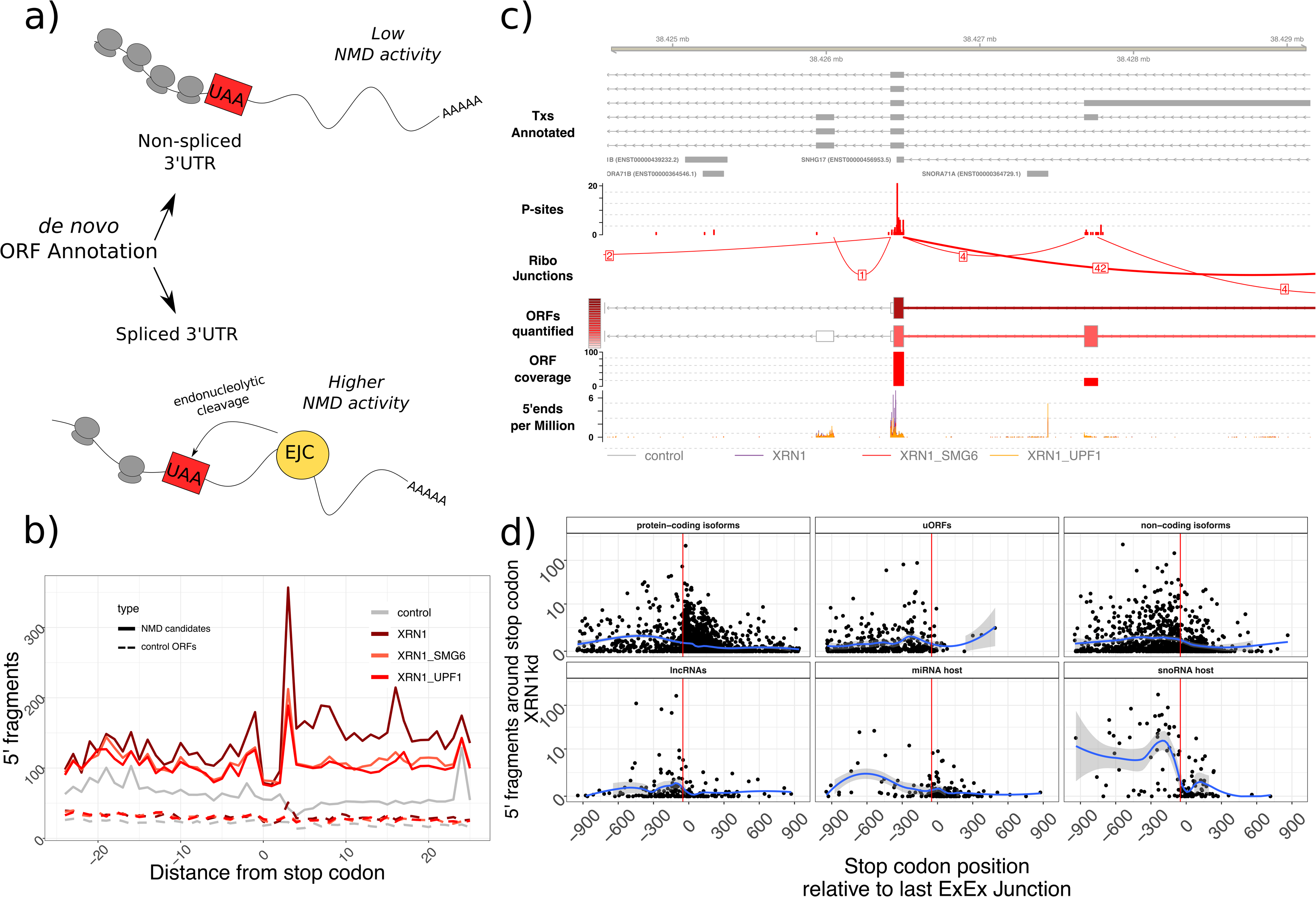
*De-novo* annotation of NMD candidates. a) Schematic annotation of NMD candidates; EJC = Exon Junction Complex. b) Aggregate profiles of 5’ fragments around stop codons of NMD candidates and control ORFs from the same genes. c) Example of a (not previously annotated) translated ORF in the *SNHG17* gene. d) Number of 5’ fragments observed in an XRN1 knockdown experiment around stop codons (*y-axis*), versus the distance between stop codons and the last exon-exon junction (*x-axis*), for different transcripts/ORF classes. Smoothing was carried out by a generalized additive model (*gam* in R, with default parameters). The red vertical line indicates 50 nucleotides upstream of the last exon-exon junction.

To further explore the dependency of NMD with regards to the location of PTCs as well as the transcript type, we determined the number of endonucleolytic cuts at the stop codon as a function of PTC distance to the last exon-exon junction. We observed an increase in degradation for NMD candidate ORFs for all the surveyed ORFs (including uORFs; Figure 3d). As previously reported^25^, ORFs in snoRNA host genes (such as *SNHG17*, Figure 3c) showed the highest degradation profile, while other categories exhibited a lower amount of degradation. In summary, *ORFquant* is an efficient method to identify mature mRNAs, quantify the translation output of different transcript isoforms from the same gene, and to infer transcript-specific cytoplasmic fates.

### A subset of genes translates different major ORFs in different cell lines

To investigate the patterns of alternative ORF usage across different conditions, we ran *ORFquant* on Ribo-seq datasets from 6 different human cell lines (Supplementary Table 1, Supplementary Data 1, Figure 4a), with newly generated data for K562 and HepG2 cell lines complementing previously published libraries from HEK293, HeLa, U2OS and Jurkat cells^18,26–28^. For each dataset we observed the same trend described in Figure 1d, with most genes showing translation of one major ORF, and hundreds of genes showing sustained translation of multiple ORFs, with a weak dependency on the overall Ribo-seq signal (Supplementary Figures 5, 6). Across all cell lines, we detected ORF translation for ~17,000 genes (excluding pseudogenes), with ~89% of them annotated as protein-coding genes.

**Figure 4:**
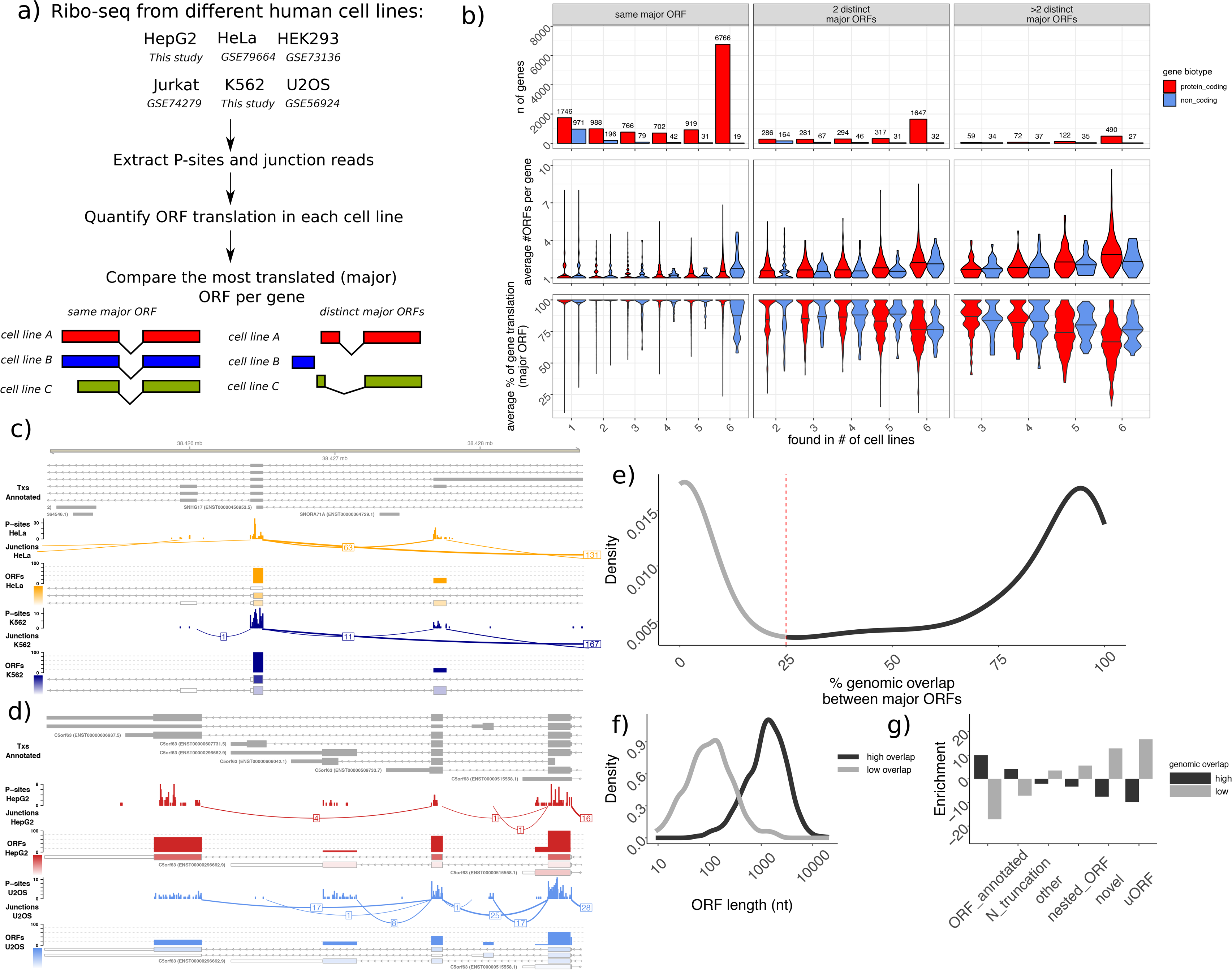
Diversity in gene translation across cell lines. a) Workflow for the analysis of the different datasets. b) Number of genes (*top*), average number of detected ORFs (*middle*) and average % of gene translation (*bottom*), for each number of cell lines where the gene harbored a detected ORF. Colors indicate the gene biotype. Genes translating one or more distinct major ORFs across cell lines are shown in different panels. The maximum width of each violin plot is the same for each panel, and the median value is shown as a black bar. c) Detected ORFs and Ribo-seq signal in the *SNHG17* gene in HeLa and K562 cells. d) Detected ORFs and Ribo-seq signal in the *C5orf63* gene in HepG2 and U2OS cells. e) Distribution of overlap between multiple major ORFs from the same gene. f) Length (in nucleotides) of major ORFs with low and high overlap. g) Enrichment of different categories for major ORFs with high and low degree of overlap.

For each gene and cell line, we defined the major ORF as the most translated ORF from a gene, regardless of its positional features and existing ORF annotation. For ~77% of the quantified genes, the same ORF was consistently identified as the major translated ORF in all the assayed cell lines (Figure 4b). For ORFs in non-coding RNAs, we detect a more cell-specific pattern of major ORF usage. However, a few dozen non-coding genes displayed translation of the same major ORF: one such example is again *SNHG17*, where the translation of an ORF terminating at a PTC (Figure 3c) is consistently detected across the assayed cell lines (Figure 4c).

As expected, genes translated in all cell lines showed overall higher Ribo-seq signal. However, we did not observe a clear dependence between number of distinct major ORFs across cell lines and overall gene translation (Supplementary Figure 7). Two or more distinct major ORFs were identified in 18% and 5% of genes, representing candidate major ORF switching events across cell lines (Figure 4b). At a closer look, we observed that genes translating multiple major ORFs also displayed a more complex mixture of translated ORFs. Consequently, translation of the major ORF for those genes accounted for a lower percentage of total gene translation (Figure 4b, lower panel).

ORF diversity is created by different mechanisms: differences in alternative splicing of internal coding exons (Supplementary Figure 8), alternative transcriptional start sites (Supplementary Figure 9), or alternative usage of last exons (Figure 4d). Genes exhibiting translation of multiple major ORFs showed an enrichment for GO categories like GTPase regulator (Supplementary Figure 10), a category also enriched in genes expressing multiple major transcript isoforms across human tissues^29^. However, in ~40% of the cases, distinct major ORFs translated across cell lines showed a low degree of overlap (Figure 4e) despite coming from the same genes, i.e., largely unrelated to differences in local alternative splicing events. This low overlap reflected the presence of alternative usage of uORFs or other small ORFs (Figure 4f,g), which can represent the major translation product of a gene in specific cell lines (Supplementary Figure 11).

Taken together, these translation estimates indicate that the presence of one dominant ORF agrees across multiple cell lines for the majority of genes. For ~20% of the translated genes, however, highly translated small ORFs and/or several transcripts expressed at sustained levels create a substantial level of complexity in protein synthesis, with distinct ORFs accounting for the majority of the gene translational output in different cell lines.

### Agreement between protein abundance and synthesis estimates depends on proteome coverage and transcriptome complexity

Ribo-seq reflects the density of elongation-competent 80S ribosome, and thus active protein synthesis, but an increased signal at a specific location may also represent stalled, inactive ribosomes. We therefore examined whether our translation quantification reflects the abundance of the synthesized protein product. Using a comprehensive custom protein database derived from the set of identified ORFs (Supplementary Figure 12, Supplementary Data 2), we estimated proteome-wide steady-state protein abundance using published deep mass spectrometry data^30,31^ for the same cell lines outlined above (Figure 4a). We detected between 7,000 and 8,000 proteins per cell line (Supplementary Figure 13, Supplementary Data 2), and performed label-free quantification using signal from unique peptides only (Methods). To estimate the ability of both techniques in quantifying protein synthesis/abundance, we divided proteins based on the number of exon or junction features covered by Ribo-seq (with >=1 read mapping, i.e. independent of the exact number of mapping reads), and by the number of detected unique peptides (irrespective of their intensity). In cases where 0 - 3 unique peptides were detected, the correlation (in log space) between HEK293 *ORFquant*-derived estimates of translation (*ORFs_pM*) and steady-state protein abundance (*iBAQ*) measured ~0.52 (Figure 5a). However, for proteins having >9 uniquely mapping peptides and >8 covered features (n >1900), the correlation between *ORFquant* estimates of translation and protein abundance reached the value of ~0.84. The same phenomenon was observed for all the assayed cell lines (Figure 5b). A clear dependency on the number of unique peptides was also observed when correlating *iBAQ* values with transcript abundance estimates from RNA-seq, albeit with lower correlations (Supplementary Figure 14).

**Figure 5:**
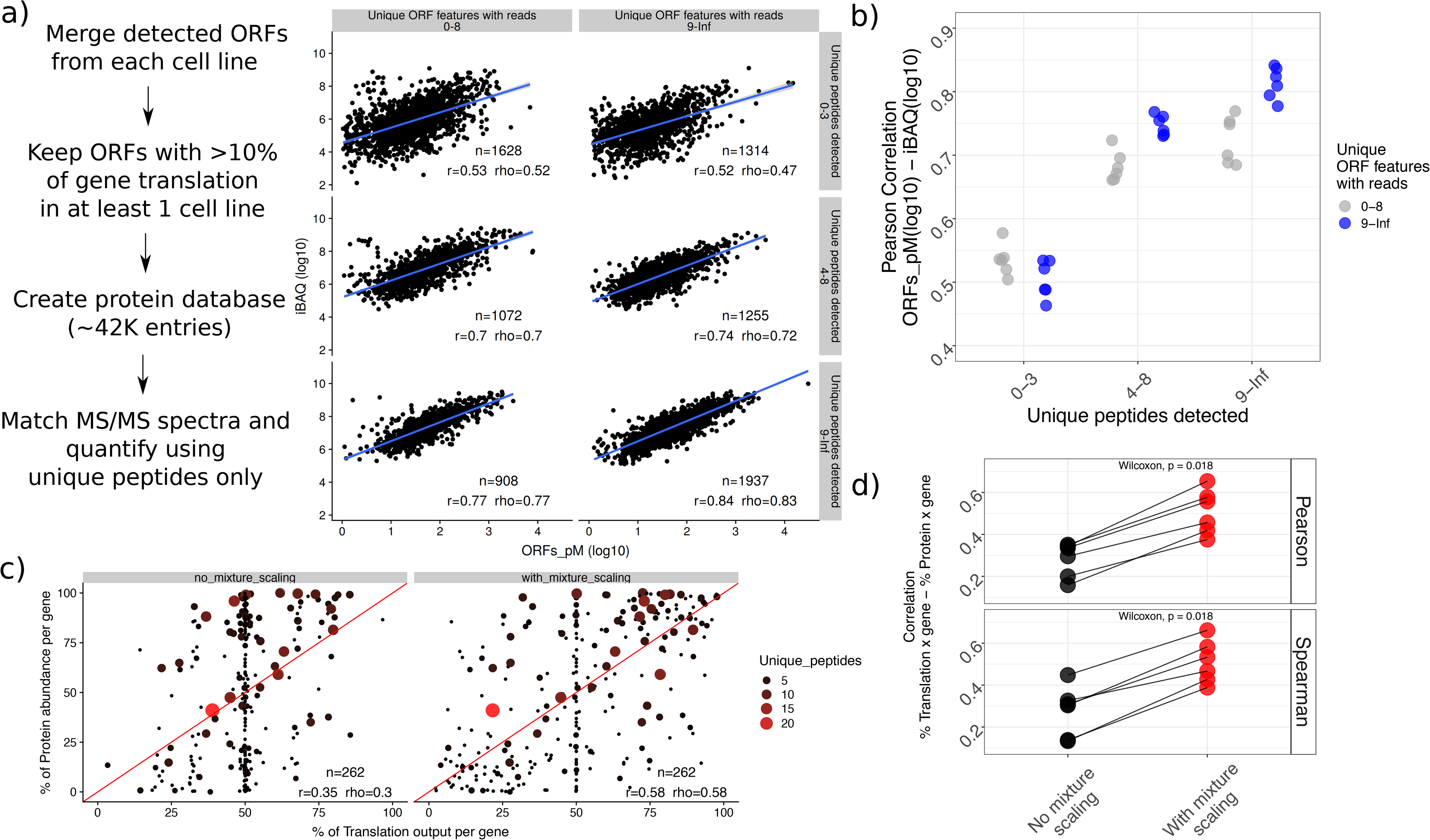
Agreement of protein synthesis with steady-state protein abundance estimates. a) Workflow of the proteomics analysis (*left*). On the right, iBAQ values (*y-axis*) versus length-normalized translation quantification estimates (*ORFs_pM*); proteins are split in multiple groups based on the number of detected unique peptides (from proteomics) or unique covered features (from Ribo-seq). b) Correlation values (as in a) shown for all the assayed cell lines. c) % of protein abundance per gene (*y-axis*) plotted against % of gene translation (*x-axis*), with translation quantification performed with (*right*) and without (*left*) adjusting for the presence of multiple ORFs per gene. Size and color of each data point indicate the number of unique peptides detected. d) Correlations from c) shown for the 6 cell lines. P-values derived from paired Wilcoxon rank-sum test (one-sided).

When comparing the fraction of total gene translation to the fraction of total gene protein abundance for the few dozen genes with mass spectrometry matches to multiple detected protein isoforms, we observed a correlation of 0.58 (Figure 5c). Here, only few proteins harbored >9 uniquely mapping peptides, thus limiting our ability in reliably estimating their abundance (Figure 5a). We observed lower correlations when skipping the ORF-specific scaling step during translation quantification, highlighting the importance of accounting for the presence of multiple translated ORFs per gene (Figure 5c). The same pattern was observed for the other cell lines analyzed (Figure 5d, Supplementary Figure 15). A slight increase in correlations was detected when using all Ribo-seq reads (instead of uniquely mapping reads only) to derive translation estimates (Supplementary Figure 16), likely resulting from a better quantification in repetitive regions.

Taken together, these results show excellent correlations between *ORFquant* quantification and steady-state protein abundance, subject to the limitations in coverage that leads to lower agreement between Ribo-seq and shotgun proteomics.

## Discussion

Only a fraction of known, annotated transcript structures are expressed in a specific context, and only a fraction of those structures are exported to the cytoplasm and eventually translated into functional proteins. This observation inspired us to devise a simple strategy to identify the subset of translated ORFs across transcript isoforms from Ribo-seq data, by discarding a substantial fraction of transcript structures with no support. The marked nuclear localization of annotated but discarded RNAs (Figure 2a) indicates that these structures are not present at translating ribosomes, i.e. that they are not expressed in the assayed condition or that they represent pre-mRNA intermediates which are either rapidly degraded or retained in the nucleus. Our strategy therefore resulted in a markedly improved mapping of Ribo-seq exonic and junction reads to their possible transcripts of origin (Figure 1d), allowing for ORF-specific translation estimates.

The quantification of transcript isoform expression is a well-studied problem in RNA-seq, with popular methods applying iterative methods (such as the expectation-maximization algorithm) to resolve the mixture resulting from multiple transcripts^19,32,33^. However, resolving the mixture of multiple transcript isoforms can be challenging for some genes, especially in the absence of coverage on unique transcript features. We could show how a top performing algorithm designed to solve this problem with high accuracy displays high variability in its estimates for such cases (Supplementary Figure 2), illustrating how short-read sequencing data, such as resulting from Ribo-seq, does not provide sufficient information to impeccably quantify translation for each and every gene. The rapidly increasing availability of full-length transcript sequence data based on long-read sequencing^34^ holds great promise in solving these complex scenarios.

While polysome profiling experiments (Figure 2d) and label-free quantification of the protein product (Figure 5b) support the Ribo-seq-based estimates of relative ORF translation levels, we believe that additional efforts can improve ORF-specific quantification of translation. A more accurate approach will have to address the issue of variable Ribo-seq coverage along the ORFs, which reflects the complex dynamics of translation. However, the impact of different features, such as experimental biases, codon composition or RNA structural features^35^, on Ribo-seq coverage remain to be understood. Our approach also uses a strict definition of ORFs that requires a canonical start codon and does not account for overlapping frames. It is still an open question how to correctly define the precise boundaries of translated elements that account for non-canonical start codons and signals from overlapping frames, such as from upstream ORFs^36^ or complex gene structures in compact genomes such as found in viruses and organelles. Our strategy enabled us to detect thousands of lowly translated ORFs in transcript isoforms of protein coding genes that are annotated as non-coding, consistent with current models for mRNA surveillance such as NMD (Figure 3). Similarly, we observed that many detected ORFs in non-coding RNAs show high degradation profiles at their stop codons, especially pronounced in snoRNA host genes (Figure 3d). This well-known phenomenon is therefore important to consider when addressing the protein-coding ability of transcripts based on ribosome occupancy. In turn, the ability to identify NMD target candidates can provide an advantageous starting point for further research into defining the features of co-translational mRNA surveillance and its links to protein quality control^37^.

Expanding our analysis across multiple cell lines allowed us to assess the complexity of translation per gene for both coding and non-coding genes (Figure 4b). We found the majority of genes to be translating the same major ORF (including highly translated ORFs in non-coding RNAs, Figure 4c), but we also detected distinct ORFs used for the major translation product in different cell lines in thousands of genes. These genes showed an overall more complex pattern of transcript expression, with sustained translation of many transcripts, and thus posing a difficulty in defining clear isoform switching event. In this context, the presence of highly translated small ORFs in protein-coding genes (Supplementary Figure 11), which may play gene regulatory roles rather than expand the proteome, adds further complexity. Unfortunately, the limited amount of data at hand (often lacking replicate information) and the heterogeneity of protocols adopted by different labs, pose challenges to precisely quantify the contribution of different mechanisms in promoting diversity (or lack thereof) in protein synthesis.

Despite potential limitations, we observed a substantial agreement between our estimates of translation and steady-state protein abundance. The level of agreement between mRNAs and proteins has been subject to intense debate^38^; our results indicate that for thousands of genes, shotgun proteomics experiments and sequencing of ribosome-occupied RNA fragments do show excellent agreement, albeit with expected dependencies on the reliability with which we can quantify the levels of translation and protein abundance (Figure 5a). An increasing availability of Ribo-seq and proteomics data in a single controlled environment will improve our understanding of this relationship and help to pinpoint interesting cases in which this correlation deviates from expectation. While our analyses provide a promising starting point for the investigation of transcript-specific protein production, the current scarcity of matching data specifically limits our ability to validate the translation of alternative protein isoforms per gene. A recent study demonstrated how protein isoforms engage with distinct protein-protein interaction networks, with such interactions being as different as the ones involving proteins from distinct genes^39^. With both proteomics^40,41^ and transcriptomics^42^ techniques rapidly advancing at a fast pace, our study demonstrates the unique advantage of ribosome profiling in characterizing and quantifying cytoplasmic gene expression programs, at the interface between RNA and protein.

## Methods

### *ORFquant* - Transcript/ORF filtering

Gene models from the GTF annotation are flattened to obtain coordinates about exonic bins or junctions, together with the set of transcripts they map to. Next, P-sites positions and junction reads (from all read lengths) are mapped to such features, to obtain *positive* (with at least one read count) or *negative* features (with no reads). Internal features are then defined as features contained between the coordinates of the first (most upstream) and last (most downstream) positive features.

The filtering procedure is then applied: initially, an empty vector of positive features is created, and such a vector is updated at each step, adding (when present) new positive features contained by the analyzed transcript. After creating the empty vector, the set of annotated transcripts is analyzed, applying the following rules for each transcript *Tx*_*i*_:

1. *Tx*_*i*_ contains a novel positive feature: *Tx*_*i*_ is selected and each previously selected *Tx*_*j*_ is re-analyzed: If all the positive features of *Tx*_*j*_ are also contained in *Tx*_*i*_, *Tx*_*j*_ is discarded.
2. *Tx*_*i*_ does not contain a novel positive feature: *Tx*_*i*_ is initially selected, but it is compared with each previously selected structure *Tx*_*j*_. Two possible scenarios are evaluated:

i. All the positive features of *Tx*_*i*_ are also contained in *Tx*_*j*_: if *Tx*_*j*_ has more positive features than *Tx*_*i*_, or fewer negative *internal* features than *Tx*_*i*_, *Tx*_*i*_ is discarded
ii. All the positive features of *Tx*_*j*_ are also contained in *Tx*_*i*_: if *Tx*_*i*_ has fewer negative *internal* features than *Tx*_*j*_, *Tx*_*j*_ is discarded.

This greedy strategy reduces the number of transcripts that is necessary to cover all the positive features (features with reads), trying to minimize the presence of negative features (features with no reads). We select ORFs following the same rules, this time using exonic bins and splice junctions derived from the ORF structures.

### *ORFquant* - ORF finding

As in the RiboTaper^18^ method, only ATG is considered as potential start codon, and the p-value for the multitaper method applied to the candidate ORF P-sites track must be below 0.05. To select ORFs with in-frame P-sites and account for local off-frame effects, we require the average signal on each covered codon to be >50% in frame. The same strategy is used to select the start codon for each ORF, requiring >50% average in-frame codon signal between each candidate ATG and the next.

### *ORFquant* - ORF quantification

After the ORF finding step, ORF filtering and quantification is subsequently performed, using the length-normalized Ribo-seq coverage *Cov* on each ORF feature.

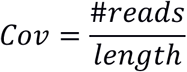

P-sites positions are used to calculate coverage on exonic regions, while spliced reads for junctions. Length is set to 60nt for junctions, according to the possible nucleotide space covered by a spliced read of ~30nt.

A feature *F* can be unique to one ORF or shared between multiple ORFs. For each ORF, we calculate the average coverage on unique features *AUcovUN*, using the coverage *Cov*_*Fu*_ on each of the unique features *F*_*u*_.

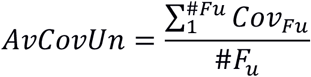

The same calculation is performed for all features *Fall* mapping to the ORF.

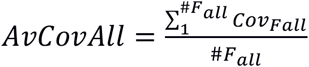

A scaling factor *C*_*ORF*_ (with a minimum value of 0 and a maximum of 1) is calculated, for each ORF, using the ratio between *AUCovUN* and *AUCovAll*. Such scaling factor represents the fraction of Ribo-seq signal that can be attributed to the ORF.

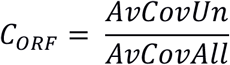

When no unique feature is present in one ORF (all regions are shared with other ORFs), the signal at each feature is adjusted using the quantification performed on other ORFs, as follows: the coverage *AdjCov*_*F*_ on each feature *F* attributed to that ORF is calculated subtracting the expected signal coming from other ORFs (*ORF*_*OverF*_) overlapping that feature, using their scaling factors. In such cases, the calculation of the adjusted coverage for each feature *Fadj* is as follows:

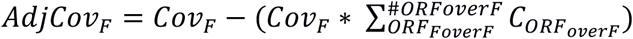

After calculating the adjusted coverage for each feature, the average of such coverage values is calculated.

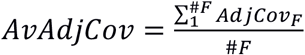

The final scaling factor is here defined by the ratio of the adjusted coverage (coverage belonging to the ORF) to the total coverage (coverage coming from all ORFs).

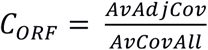

If no unique region is present in any detected ORF in the gene (all regions are shared among ORFs and no *C*_*ORF*_ value can be initially calculated), the scaling factor is derived assuming uniform Ribo-seq coverage on each ORF. The shared coverage *ShCov*_*F*_ is now simply calculated dividing it by the number of *ORF*_*overF*_ mapping to the feature *F*.

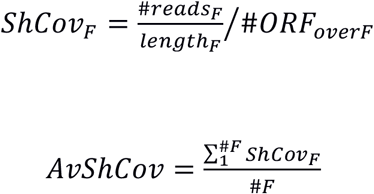

The scaling factor is again derived dividing the average shared coverage (attributed to the ORF) to total average coverage.

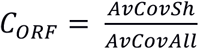

After the calculation of *C*_*ORF*_, the adjusted number of P-sites for each ORF (*P*_*ORF*_) is calculated using the raw number of P-sites mapping to the ORF multiplied by the scaling factor, to obtain ORF-specific quantification estimates.

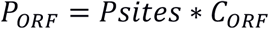

For each ORF of length *L*_*ORF*_, the scaled numbers of P-sites *P*_*ORF*_ is normalized over the entire set of detected ORFs to obtain TPM-like values, named ORFs per Million (*ORFs_pM*), using this formula:

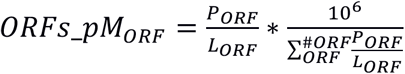

Moreover, we calculated the contribution of each ORF to the overall translation output of a single gene. Such metric, named *ORF_pct_P_sites* (or percentage of gene translation), is calculated dividing *P*_*ORF*_ by the sum of *P*_*ORF*_ of all ORFs (#*ORFg*) detected in a gene.

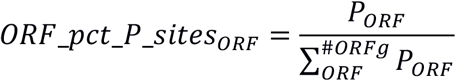

Normalization by length is here not applied, as this metric wants to quantify the amount of translation per gene coming from each ORF. The *ORF_pct_P_sites_pN* metric indicates length-normalized *ORF_pct_P_sites* values (e.g. they can be high for a short highly translated ORF).

After quantification, ORFs are subjected to a filtering step and quantification is performed again, until all ORFs are being retained.

### *ORFquant* parameters

For all cell lines, *ORFquant* was run using a cutoff of 2% of total gene translation and using only uniquely mapping reads.

#### RSEM quantification

RSEM 1.3.1 was run in strand-specific mode on Ribo-seq data using a seed length of 20, using Bowtie2 as aligner, and enabling the calculation of confidence intervals together with posterior mean estimates. *ORFquant*-derived ORF positions were used to specify the transcript sequences to use as reference. When possible, additional 15 nucleotides were added to start and end coordinates to allow for the mapping of Ribo-seq reads. The “TPM_coefficient_of_quartile_variation” column of the RSEM output was used as a proxy to monitor the variability in RSEM quantification estimates.

#### Ribosome profiling

Ribo-seq was performed as described previously^18^ and adapted for HepG2 and K562 cell lines. 5×10^6 K562 suspension cells and two 80% confluent 10 cm TC dishes of adherent HepG2 cells (DSMZ #ACC-10 and #ACC-180, respectively) were used.

Adherent cells were washed with ice-cold PBS supplemented with 100 ug/ml cycloheximide (Sigma Aldrich #C4859) and immediately snap-frozen by immersing the dishes in liquid nitrogen. The dishes were then transferred to wet ice and 400 ul of lysis buffer (1X polysome buffer (20 mM Tris-Cl pH 7.4, 150 mM NaCl, 5 mM MgCl2, with 1 mM DTT (Sigma Aldrich #43816) and 100 ug/ml cycloheximide added freshly; keep on ice), 1% (v/v) Triton X-100 (Calbiochem #648466), 25 U/ml TURBO DNase (Life Tech. #AM2238)) was immediately dripped onto the frozen cells. The cells and buffer were then scraped off and left to thaw on one side of the dish, mixing them using a pipet tip.

Suspension cells were supplemented with 100 ug/ml cycloheximide, pelleted for 5 min at 300 g and washed with ice-cold PBS + 100 ug/ml cycloheximide. The washed cell pellet was immediately snap-frozen in liquid nitrogen. 400 ul of ice-cold lysis buffer was added, and the cells were put on wet ice to thaw, mixing them using a pipet tip.

The cells were left to lyse for 10 min on ice, followed by 10x trituration through a 26-G needle. After centrifugation for 10 min at 20‘000g at 4°C the clarified supernatant was transferred to a pre-cooled tube on ice. For nuclease footprinting, 400 ul of lysate were supplemented with 1000 U of RNase I (Life Tech. #AM2295) and incubated in a thermomixer set to 23°C, shaking at 500 rpm for 45 min. Footprinting was stopped by adding 260 U of SUPERASE-In (Life Tech. #2696).

To recover ribosomes two MicroSpin S-400 HR columns (GE Healthcare #27-5140-01) per 400 ul of sample were equilibrated with a total of 3 ml of polysome buffer. The columns were drained by spinning for 4 min at 600 g, then the sample was applied and spun for 2 min at 600g. Three volumes of Trizol LS (Life Tech. #10296010) were added to the flow-through and RNA was extracted using the Direct-zol RNA Mini-Prep kit (Zymo Research #R2052) as per the manufacturer’s instructions. RNA was quantified using the Qubit RNA Broad Range Assay (Life Tech. #Q10211).

Ribosomal RNA was removed from 10 ug of footprinted RNA using the RiboZero Magnetic Gold kit (Illumina #MRZG12324) as per the manufacturer’s instructions. Footprinted RNA was precipitated from the supernatant (90 ul) using 1.5 ul of glycoblue (Life Tech. #9515), 9 ul of 3 M sodium acetate and 300 ul of ethanol by incubation for 1h at −80°C and pelleted for 30 min at max. speed at 4°C. The RNA pellet was dissolved in 10 ul of RNase-free water.

To recover the ribosome protected RNA fragments the sample was loaded onto two lanes of a 1 mm 17.5% Urea-PAGE gel along with 27 nt and 30 nt RNA markers. The gel was run in 1X TBE at 250 V for 80 min and stained for 3 min in 1X SYBR gold (Life Tech. #S11494) in 1X TBE. Sample bands between 27 nt and 30 nt were excised and crushed by spinning through a punctured tube. RNA was extracted by soaking the gel pieces in 400 ul of RNA extraction buffer (400 mM NaCl, 1 mM EDTA, 0.25% (wt/v) SDS) for 2 h, rotating at room temperature. The supernatant was supplemented with 1.5 ul of glycoblue and 500 ul of isopropanol and incubated on dry ice for 30 min, followed by pelleting of the RNA for 30 min at 20‘000 g at 4°C. The pellet was dissolved in 40 ul of water.

To prepare the RNA sample for use in a smallRNA library preparation kit the sample was phosphorylated using 5 ul of 10X T4 PNK buffer and 1 ul of T4 PNK (NEB #M0201), 1 ul of SUPERASE-In, 2.5 ul of 10 mM ATP and 0.5 ul of 1% Triton X-100. After incubation for 1 h at 37°C RNA was precipitated and pelleted by adding 41 ul of water, 1.5 ul of glycoblue, 8 ul of 5M NaCl and 150 ul of isopropanol as described before. Libraries were prepared using the NEXTflex Small RNA-Seq Kit v3 (BiooScientific #5132-06) as per the manufacturer’s instructions and sequenced on an Illumina NextSeq500 machine with 13 libraries pooled at 1.8 pM using one High Output Kit v2 (Illumina #FC-404-2005) with 75 cycles single-end.

### Ribo-seq and RNA-seq data processing

Ribo-seq reads were stripped of their adapters using cutadapt^43^. Randomized UMI sequences (where present) were removed, and reads were collapsed. Reads aligning to rRNA, snoRNAs and tRNA sequences were removed with Bowtie2^44^. Unaligned reads were then mapped with STAR^45^ using the hg38 genome and the GENCODE 25 annotation in GTF format. For RNA-seq and Ribo-seq, a maximum of four and two mismatches was allowed, and multimapping of to up to 20 different positions was permitted. Alignments flagged as secondary alignments were filtered out, ensuring one mapping position per aligned read. P-sites positions and junction reads were extracted using Ribo-seQC^46^ with default parameters. Statistics about the different Ribo-seq libraries are available as Supplementary Data 1. *Gviz*^47^ was used to visualize data tracks and transcript annotation.

#### Polysome profiling

DEXSeq^20^ was run to detect differential exon usage between each of the polysome fraction and the cytoplasmic abundance. Transcripts were divided based on the translation levels of their translated ORF(s) and intersected with differential exons (FDR<0.01 in at least one polysome fraction). Only genes with multiple translated transcripts were used.

#### Nuclear-cytoplasmic comparison

DEXSeq^20^ was run to detect differential exon usage between the nuclear and the cytoplasmic fraction. Differential exons (FDR<0.01) were intersected with transcript structures and only exons uniquely mapping to one transcript group (e.g. discarded transcripts, selected transcripts etc…) were selected.

#### 5’end of endonucleolytic cuts

Bigwig files for the different libraries were normalized by library size. Coordinates were lifted to hg38 and overlapped with *ORFquant*-identified stop codon positions, for both NMD candidates and controls (“canonical” stop codons taken from the same genes). A window of 50 nucleotides was used to derive spatial profiles and count the number of reads mapping around stop codons in the different conditions.

#### Merging *ORFquant* result across cell lines

ORFs were considered to be distinct if they ended at different stop codons or could not be mapped to the same transcript. Enrichments for ORF categories at different level of overlap were calculated using normalized residuals from a chi-squared test. GO enrichment was performed using the *clusterProfiler*^48^ and *topGO*^49^ packages.

### Proteomics database search

Raw data was searched using MaxQuant^31^ version 1.6.0.13, using Carbamidomethyl as fixed modification, and oxidation of Methionine and acetylation at protein N-termini as variable modifications. Quantification was performed using only unique peptides. Matching between runs was enabled. We used a custom database to perform the peptide search: *ORFquant*-detected ORFs were merged in a unique database, choosing only ORFs explaining a minimum of 10% of gene translation in at least one cell line.

### Comparison between protein abundance and translation estimates

For each protein group, *iBAQ* values were summed up for each replicate. *ORFs_pM* values were summed for all ORFs mapping to each protein group. *ORF_pct_iBAQ* values were obtained by dividing each iBAQ value for the sum iBAQ values for that gene. Protein groups mapping to multiple genes were discarded. The same procedure was applied to *ORFs_pM* values, to compare protein and translation estimates for each protein isoform. Only proteins detected by Ribo-seq (or RNA-seq) and proteomics were used. Gene-level TPM values in Supplementary Figure 14 were calculated using kallisto^32^ with default parameters.

## Data availability

Ribo-seq data for HepG2 and K562 is available at GEO under the accession X. Ribo-seq datasets for other cell lines were previously published, and accessed using the accessions GSE79664 (HeLa), GSE73136 (HEK293), GSE74279 (Jurkat) and GSE56924 (U2OS); more details about the analyzed samples can be found in Supplementary Table 1. Nuclear and cytoplasmic RNA-seq was accessed at the European Nucleotide Archive using the accession PRJEB4197. TriP-seq data was downloaded from GEO using the accession GSE69352. Transcriptome-wide tracks of 5’ ends were accessed using the accession GSE57433. Proteomics data was downloaded from the PRIDE repository under accession PXD002395. The list of P-sites positions and junction reads in the cell types analyzed is available in Supplementary Data 1. The list of quantified ORFs in the different cell lines is available in Supplementary Data 1. The final protein database is available in Supplementary Data 2, together with the parameters used to perform the MaxQuant search and the set of identified peptides and proteins.

## Code availability

*ORFquant* is available at https://github.com/lcalviell/ORFquant.

## Author contributions

Initial study conceived by L.C. and U.O. L.C. ideated and implemented the *ORFquant* pipeline, with supervision from U.O. All analysis and visualization by L.C. Ribosome footprinting libraries in K562 and HepG2 performed by A.H. Manuscript was written by L.C. and U.O. with additional input by A.H.

## Acknowledgements

The authors acknowledge funding from the German Federal Ministry of Education and Research (BMBF grant 031 A538A RBC) and the German Research Foundation (DFG grant TR175). L.C. thanks Stephen Floor (UCSF) for support and feedback during the preparation of this manuscript.

## Supplementary files

Supplementary Table 1: Summary of Ribo-seq datasets analyzed in this study.

Supplementary Data 1: Archive containing all P-sites positions and junction reads (using uniquely mapping reads), together with the set of *ORFquant* identified ORFs, for each cell line.

Supplementary Data 2: Archive containing the set of identified peptides and proteins, including their Ribo-seq statistics, the parameters used for the MaxQuant run and the custom protein database.

