## Supplementary Data 2 for "Quantification of translation uncovers the functions of the alternative transcriptome": README_Suppl2.txt

This README is part of the Supplementary Data 2 for the paper:

"Quantification of translation uncovers the functions of the alternative transcriptome"

Lorenzo Calviello, Antje Hirsekorn, Uwe Ohler

Copyright (C) 2019  Lorenzo Calviello

All data here is distributed under GPL-3 WITHOUT ANY WARRANTY without even the implied warranty of MERCHANTABILITY or FITNESS FOR A PARTICULAR PURPOSE.  
See the  GNU General Public License for more details: http://www.gnu.org/licenses/

Contact:

Contents:

ORFquant_6cells_10pct_unq.fasta	fasta file used as a database for peptide search in MaxQuant
protein_ribo_results.csv	MaxQuant quantification summarized at the protein (protein groups) level, together with Ribo-seq-related statistics.
peptide_results.csv        MaxQuant quantification summarized at the peptide level.
parameters_MQsearch.txt	parameters used for the MaxQuant run
