## Supplementary Data 1 for "Quantification of translation uncovers the functions of the alternative transcriptome": README_Suppl1.txt

This README is part of the Supplementary Data 1 for the paper:

"Quantification of translation uncovers the functions of the alternative transcriptome"

Lorenzo Calviello, Antje Hirsekorn, Uwe Ohler

Copyright (C) 2019  Lorenzo Calviello

All data here is distributed under GPL-3 WITHOUT ANY WARRANTY without even the implied warranty of MERCHANTABILITY or FITNESS FOR A PARTICULAR PURPOSE.  
See the  GNU General Public License for more details: http://www.gnu.org/licenses/

Contact:

Contents:

ORFquant_inputs_reduced.RData	RData file containing P_sites positions (uniquely mapping reads) and junction reads for the analyzed datasets.
ORFquant_outputs_reduced.RData	RData file containing a reduced ORFquant output for the analyzed datasets, including ORF quantifications and transcript/genomic ORF positions.

load the files in R using the load() function after loading the ORFquant package: e.g.

input_data<-get(load("ORFquant_inputs_reduced.RData"))

Check the ORFquant package at https://github.com/lcalviell/ORFquant to get more information about ORFquant input and output files.
