## Supplementary Figures for "Quantification of translation uncovers the functions of the alternative transcriptome"

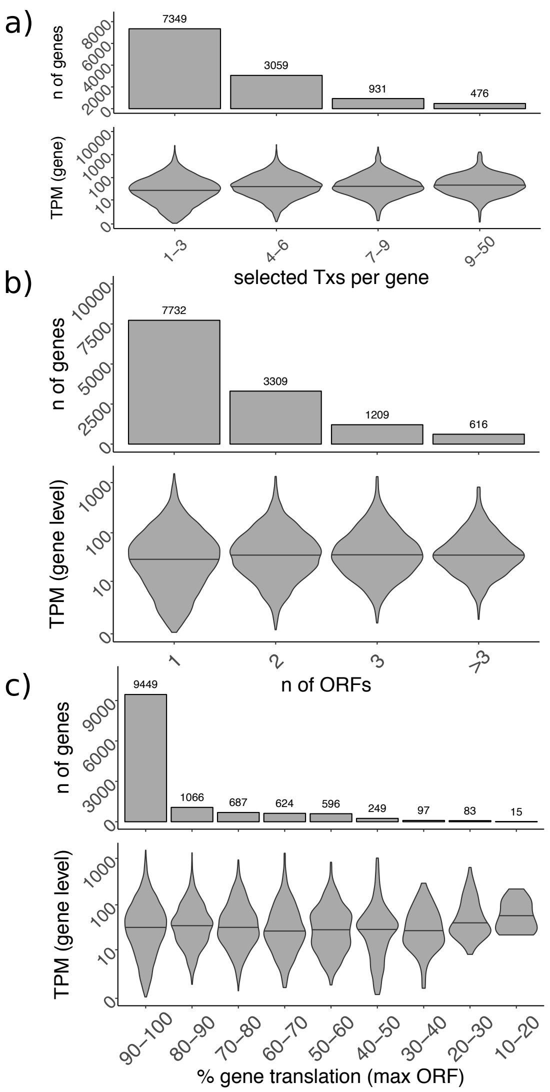


**Supplementary Figure 1:** Transcript selection and ORF quantification do not strictly depend on expression levels. In a) the number of selected transcripts per gene (x-axis) against the number of genes and their TPM levels. Number of genes and their TPM values are also plotted against b) the number of detected ORFs and c) the contribution (in percentage) of their major ORF. The maximum width of each violin plot is the same for each panel, and the median value is shown as a black bar.

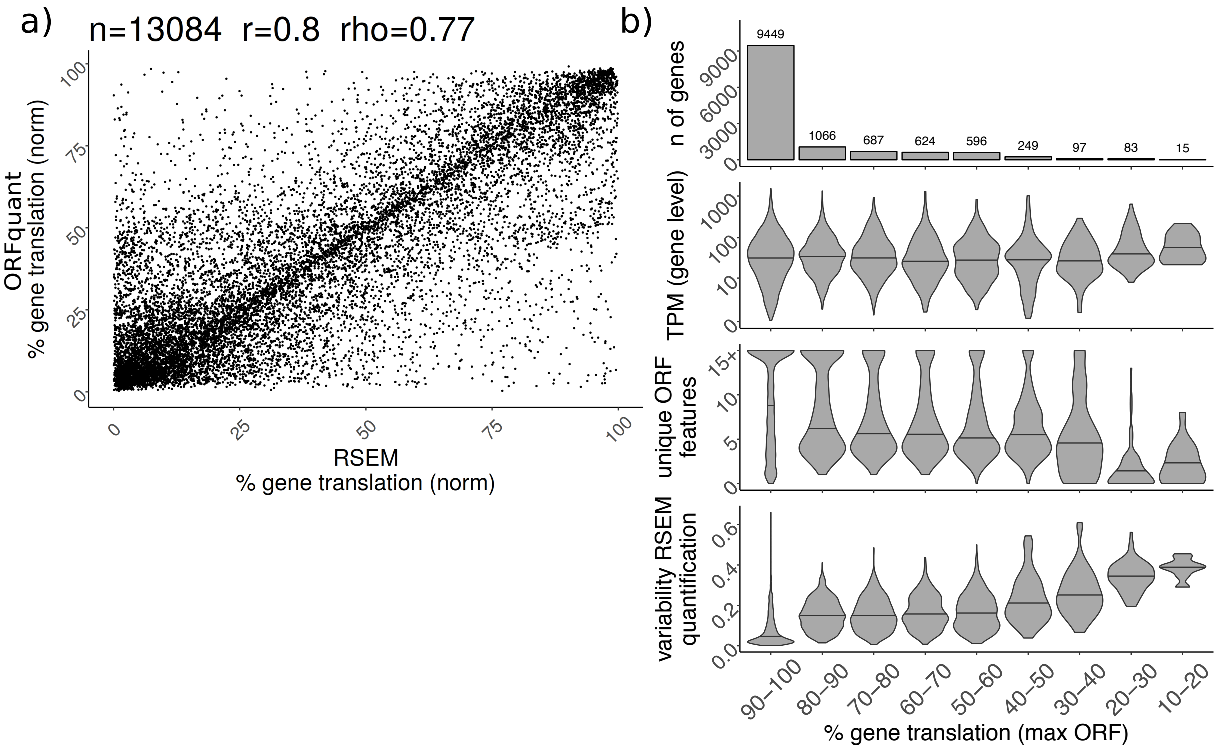


**Supplementary Figure 2:** Correlation between ORFquant and RSEM quantification estimates. In a) RSEM quantification (on the x axis, IsoPct_from_PME_TPMs) plotted against ORFquant quantification estimates (ORF_pct_P_sites_pN). Both values are calculated using length-normalized estimates. In b) the number of genes (top), their TPM values, the number of unique ORF features and the variability of RSEM quantification estimates (averaged across all the ORFs per gene) are plotted against the contribution (in percentages) of the major ORF.The maximum width of each violin plot is the same for each panel, and the median value is shown as a black bar.

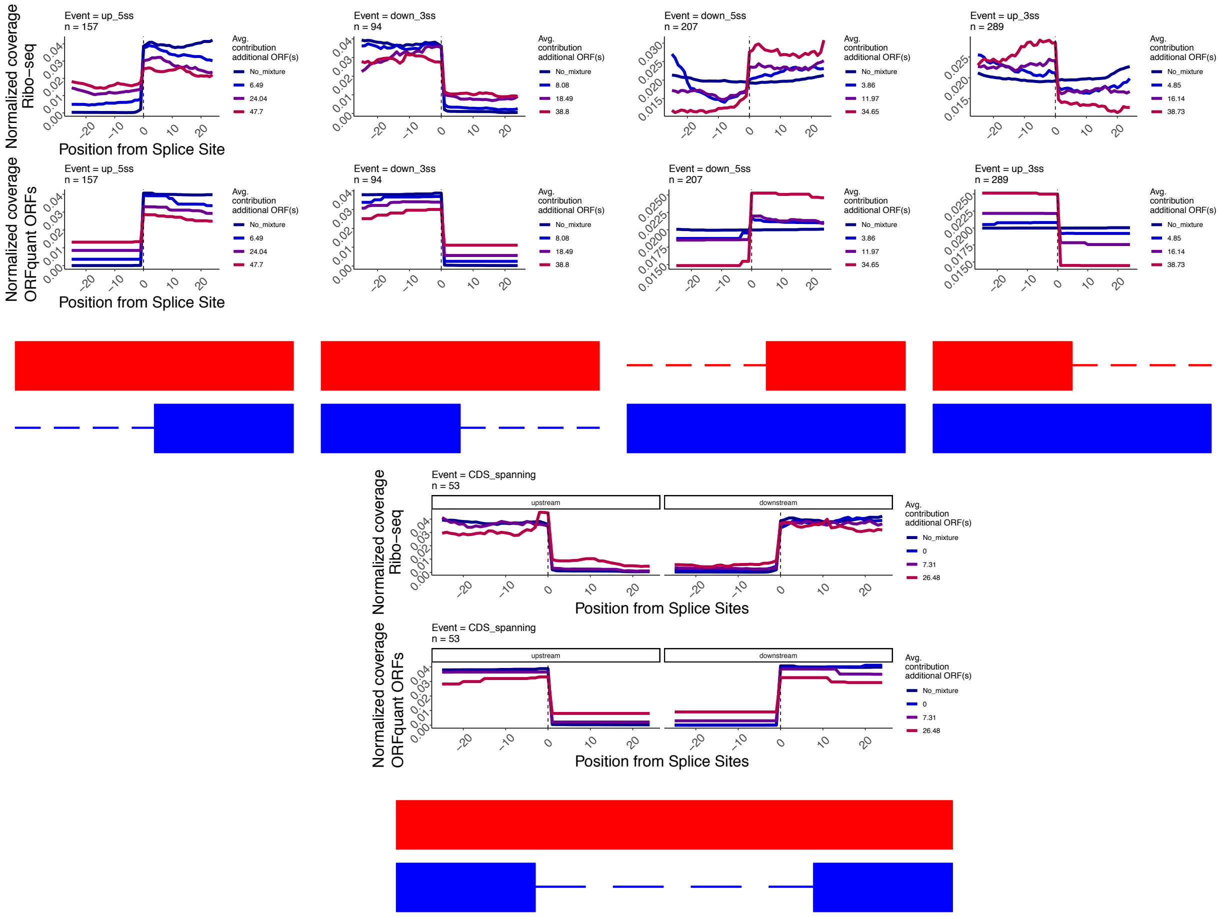


**Supplementary Figure 3:** ORFquant quantifies translation on alternative isoforms. Aggregate plots of Ribo-seq coverage (normalized 0-1 per each region) and ORF coverage (ORF_pct_P_sites_pN) over different candidate alternative splice sites. No mixture indicates the presence of a single ORF only, while other lines indicate the presence of additional ORFs, divided by their translation values. Explanatory schemes are depicted at the bottom of each plot, with blue representing the major ORF and red the additional ORF(s).

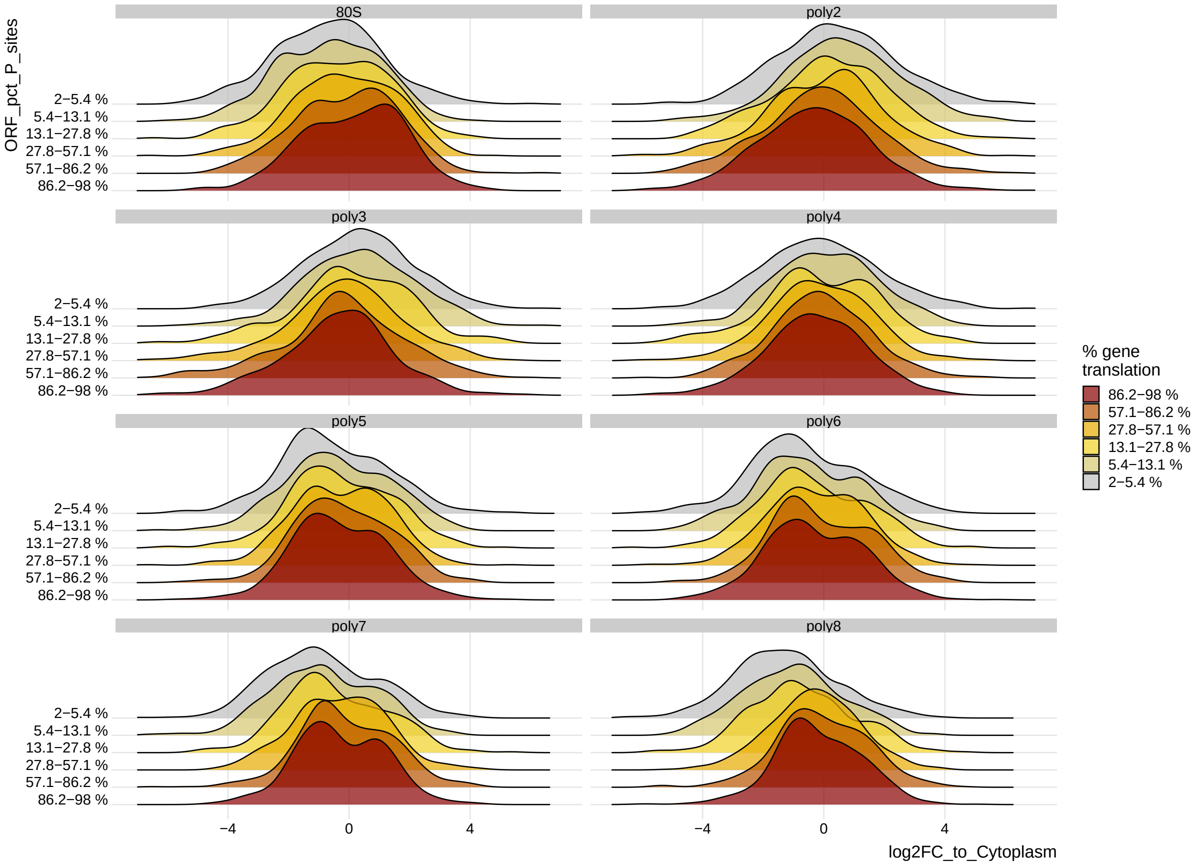


**Supplementary Figure 4:** Polysome profiles of alternative isoform with different translation output. Distributions of exonic log2 fold changes between different polysome profiles and cytoplasmic abundance. Lowly translated ORFs are depleted in heavier polysomal fractions, while highly translated ORFs show signal in all fractions.

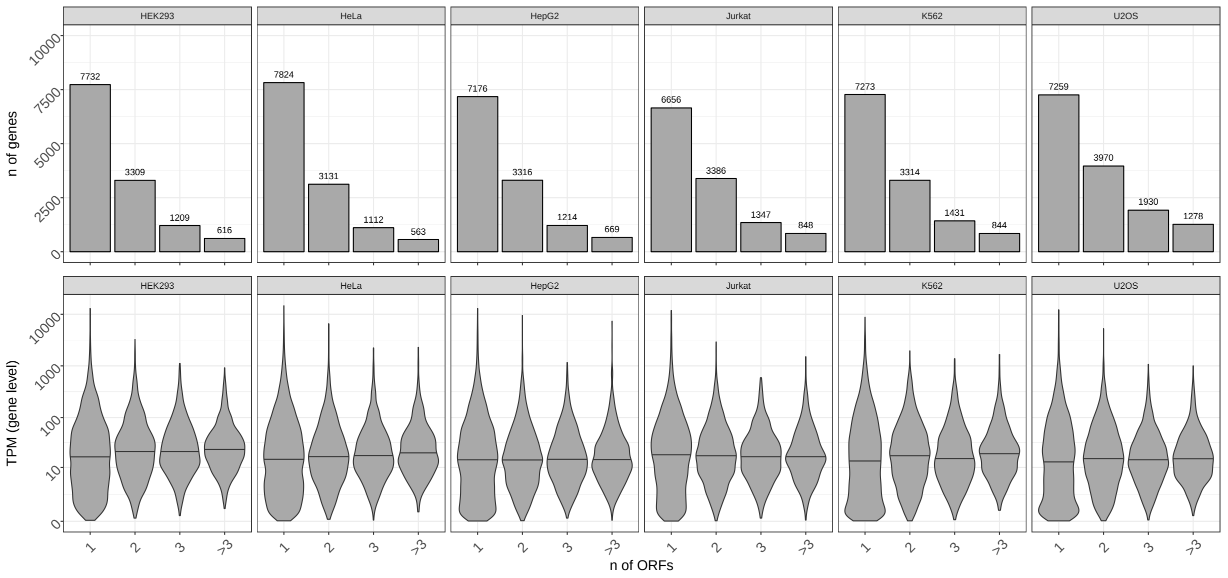


**Supplementary Figure 5:** Number of ORFs per gene detected in different cell lines. Number of genes and average Ribo-seq signal per gene (y-axes) against the number of detected ORFs (x-axis), for the assayed cell lines. The maximum width of each violin plot is the same for each panel, and the median value is shown as a black bar.

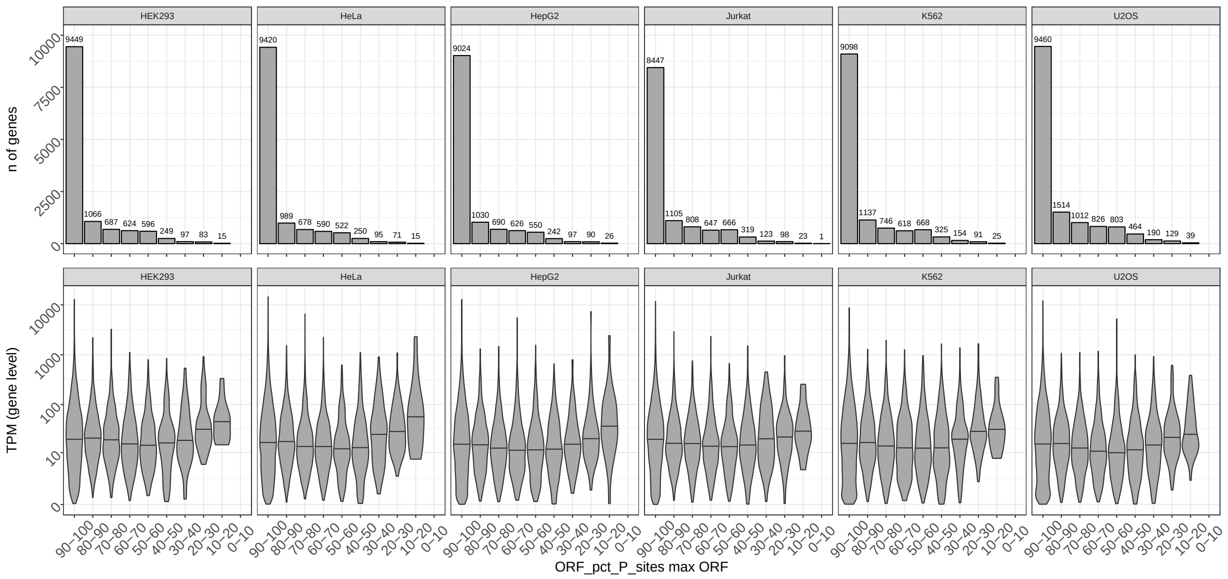

**Supplementary Figure 6:** Major ORF translation in different cell lines. Number of genes and average Ribo-seq signal per gene (y-axes) against the translation of the major ORF (x-axis), for the assayed cell lines. The maximum width of each violin plot is the same for each panel, and the median value is shown as a black bar.

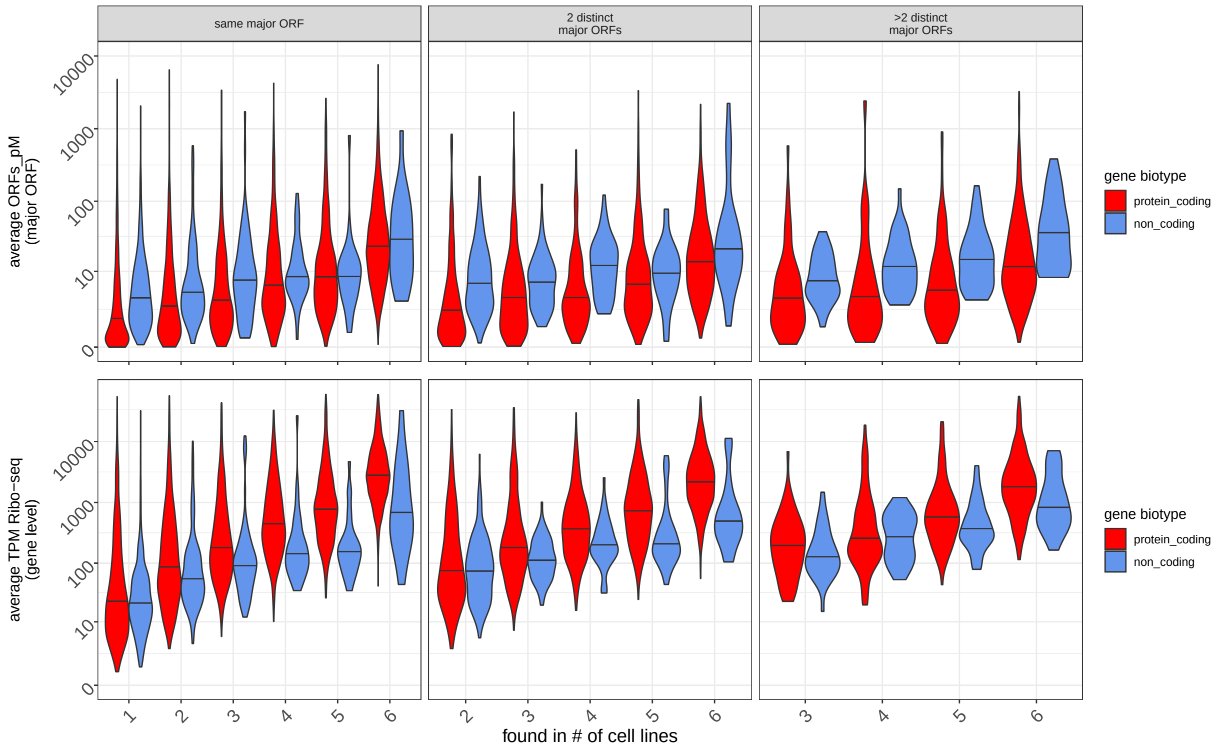


**Supplementary Figure 7:** Length-normalized translation values and overall gene signal for genes exhibiting multiple major ORFs. Average length-normalized translation of the major ORF and average Ribo-seq gene signal per gene (y-axes), plotted against the number of cell lines where the gene harbored a detected ORF. Colors indicate the gene biotype. Values were plotted dividing genes according the number of district major ORF detected across cell lines. The maximum width of each violin plot is the same for each panel, and the median value is shown as a black bar.

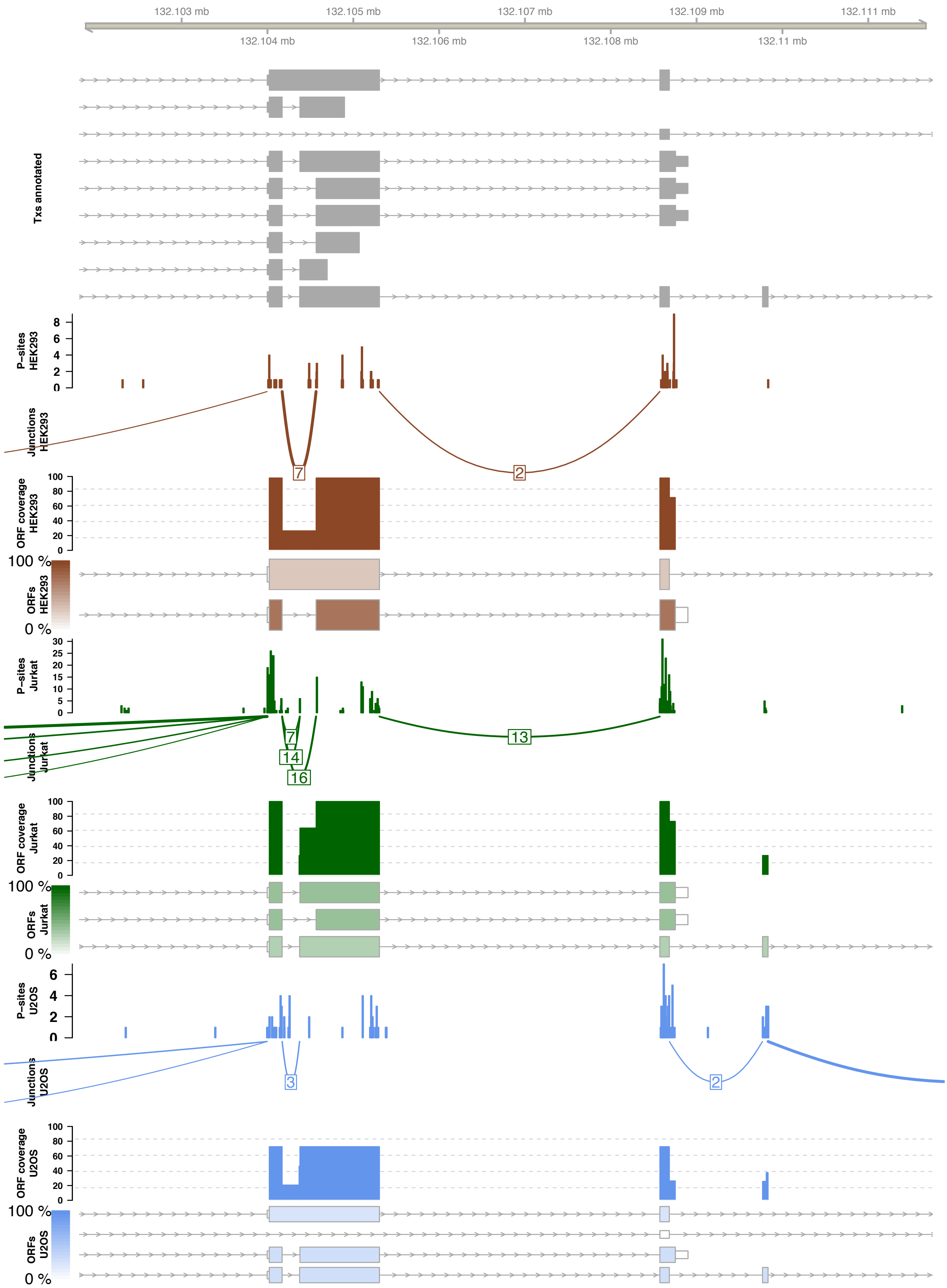


**Supplementary Figure 8:** The EP400NL gene translates multiple major ORFs across cell lines via alternative splicing.A screenshot of the EP400NL gene: displayed tracks represent, in descending order: gene annotation and (for each cell line): P-sites positions, junction reads, ORF coverage (defined as % of gene translation) and quantified ORFs.

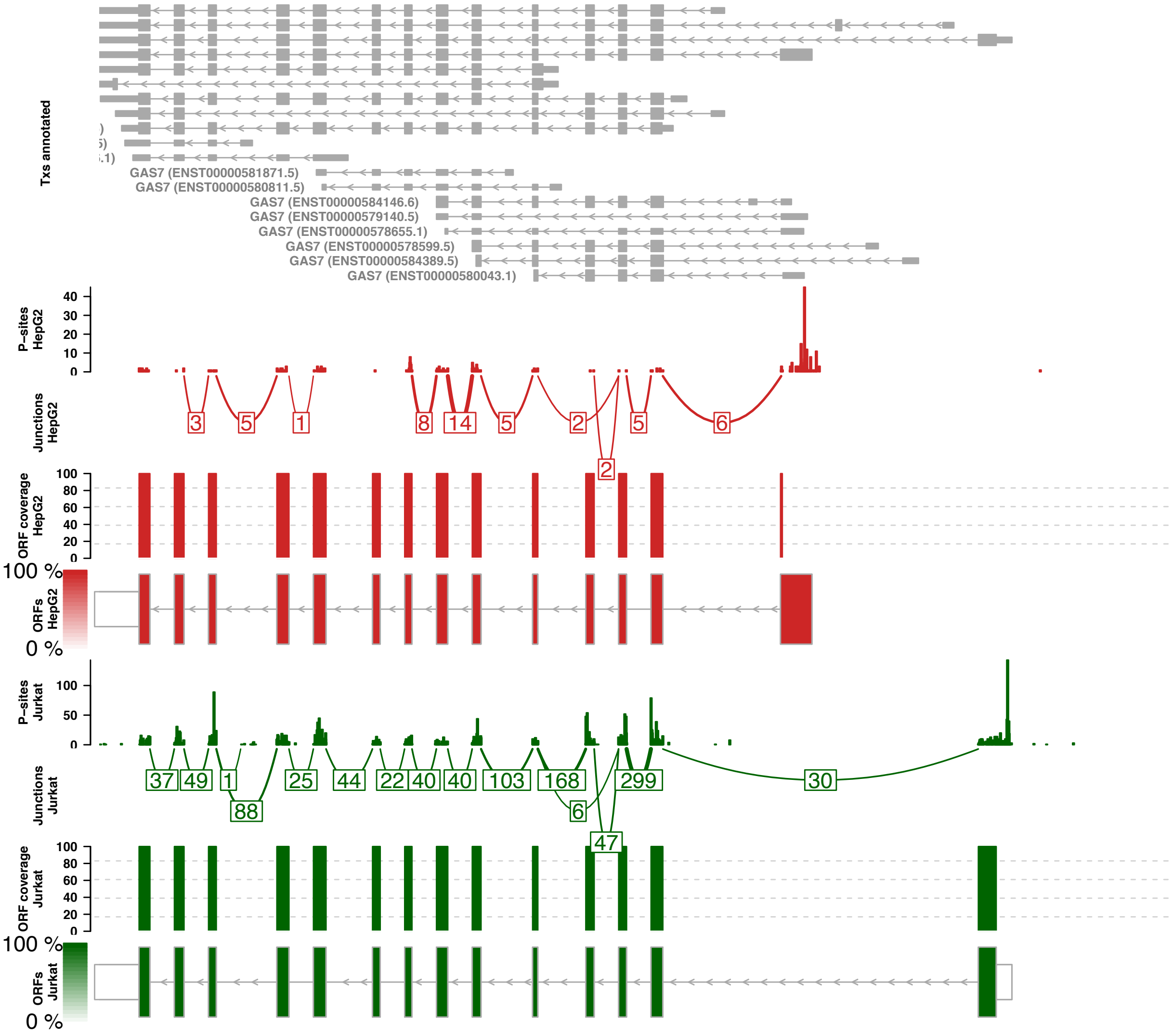


**Supplementary Figure 9:** The GAS7 gene translates multiple major ORFs across cell lines via alternative TSS usage.A screenshot of the GAS7 gene: displayed tracks represent, in descending order: gene annotation and (for each cell line): P-sites positions, junction reads, ORF coverage (defined as % of gene translation) and quantified ORFs. Introns size scaled to a maximum of 300 nt.

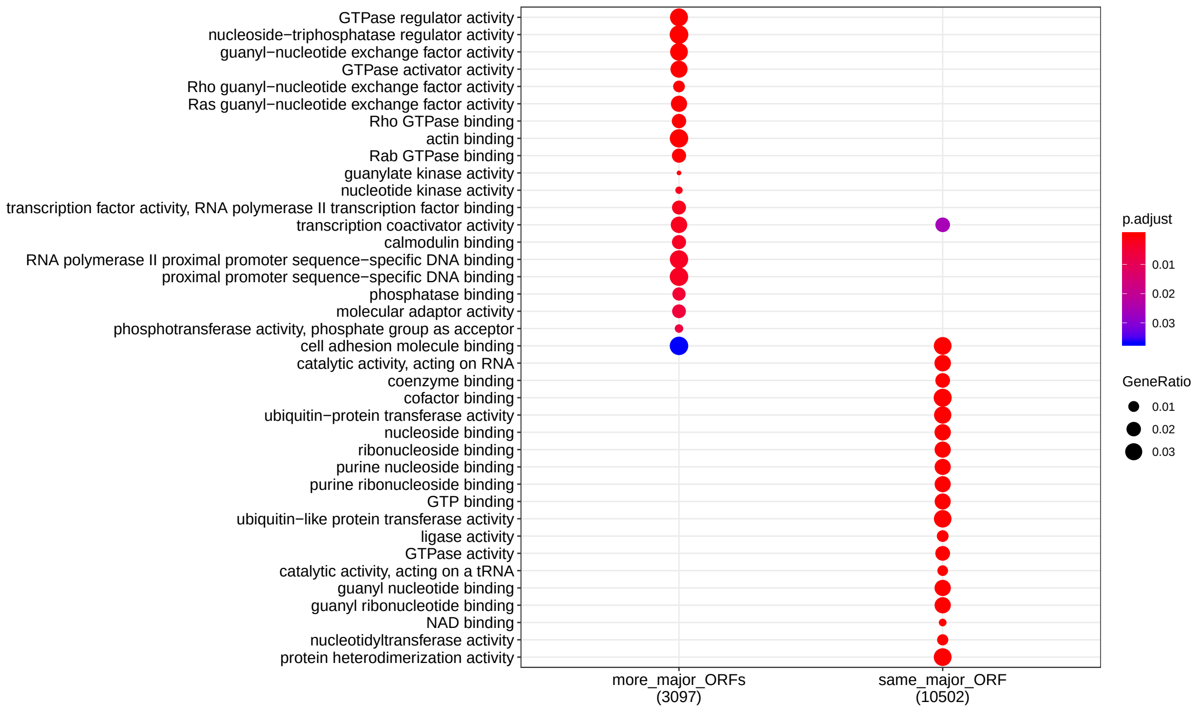


**Supplementary Figure 10:** GO analysis of single- and multi-major ORF genes. Top enriched GO categories for genes translating one (right) or multiple (left) major ORFs across cell lines.

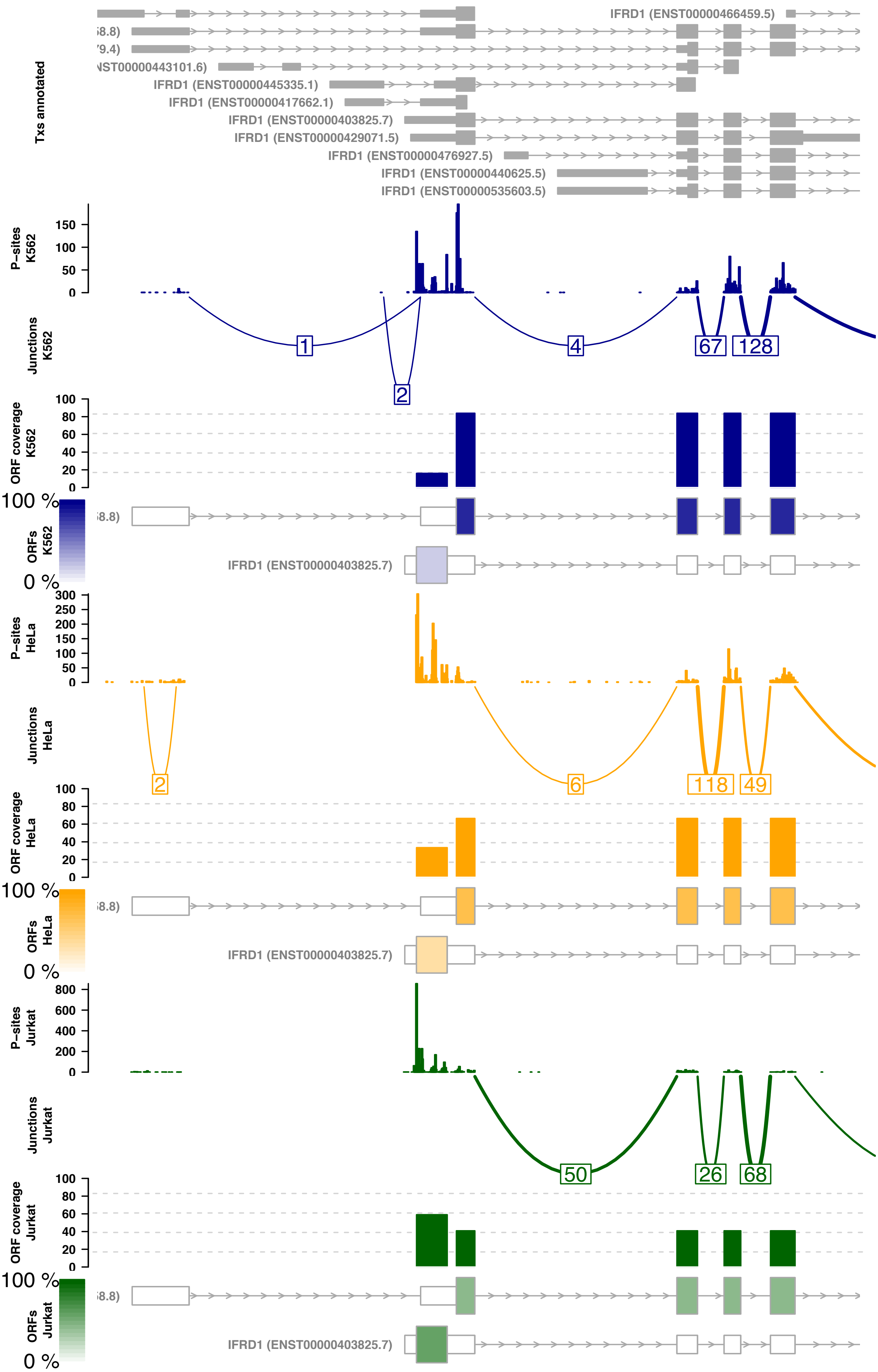


**Supplementary Figure 11:** The IFRD1 gene translates multiple major ORFs across cell lines via alternative small ORF usage. A screenshot of the IFRD1 gene: displayed tracks represent, in descending order: gene annotation and (for each cell line): P-sites positions, junction reads, ORF coverage (defined as % of gene translation) and quantified ORFs.

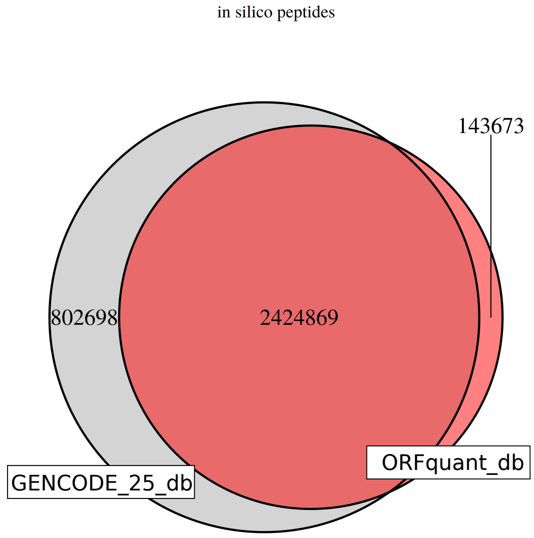


**Supplementary Figure 12:** Overlaps between in-silico generated tryptic peptides using the full GENCODE25 database or ORFquant-derived protein sequences in the 6 cell lines analyzed. Up to two missed cleavage events were allowed.

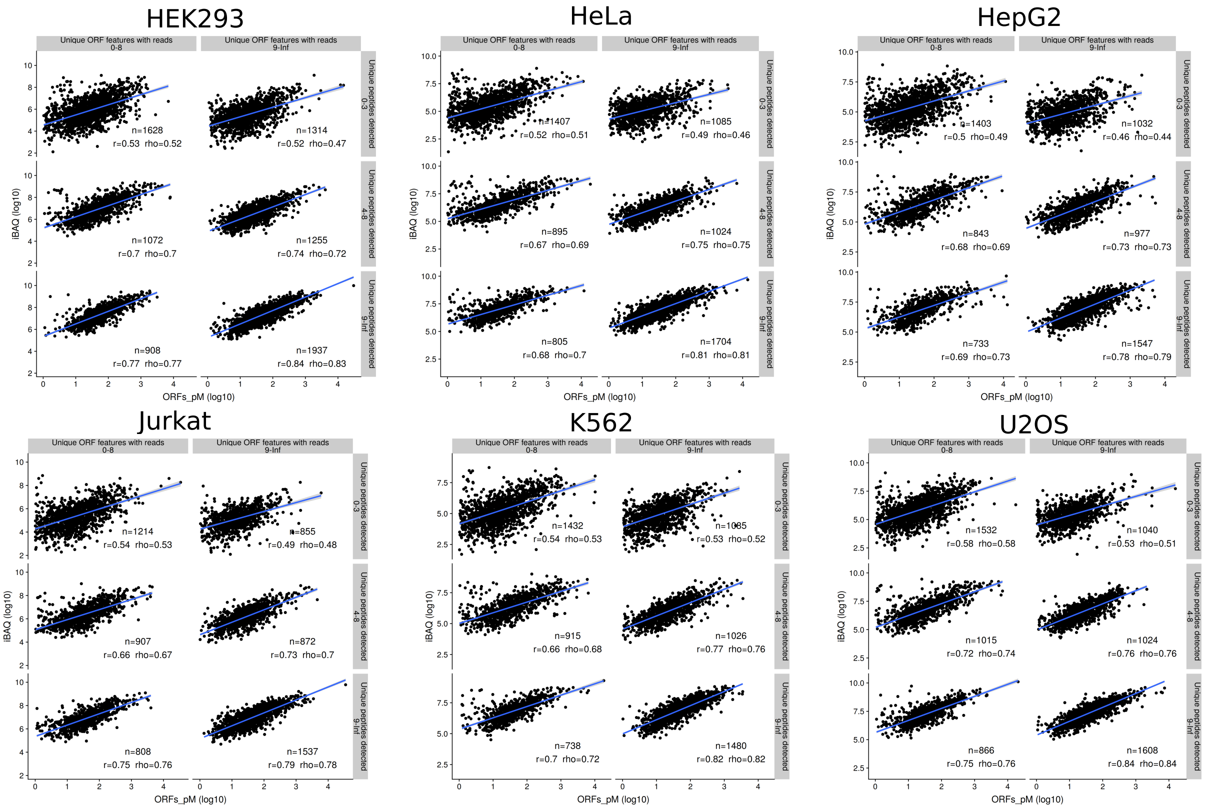


**Supplementary Figure 13:** Agreement between translation and protein abundance across cell lines. iBAQ values (y-axis) plotted against length-normalized translation quantification estimates, for each cell line. Each plot is divided by number of unique peptides (from proteomics) and unique features with reads (from Ribo-seq).

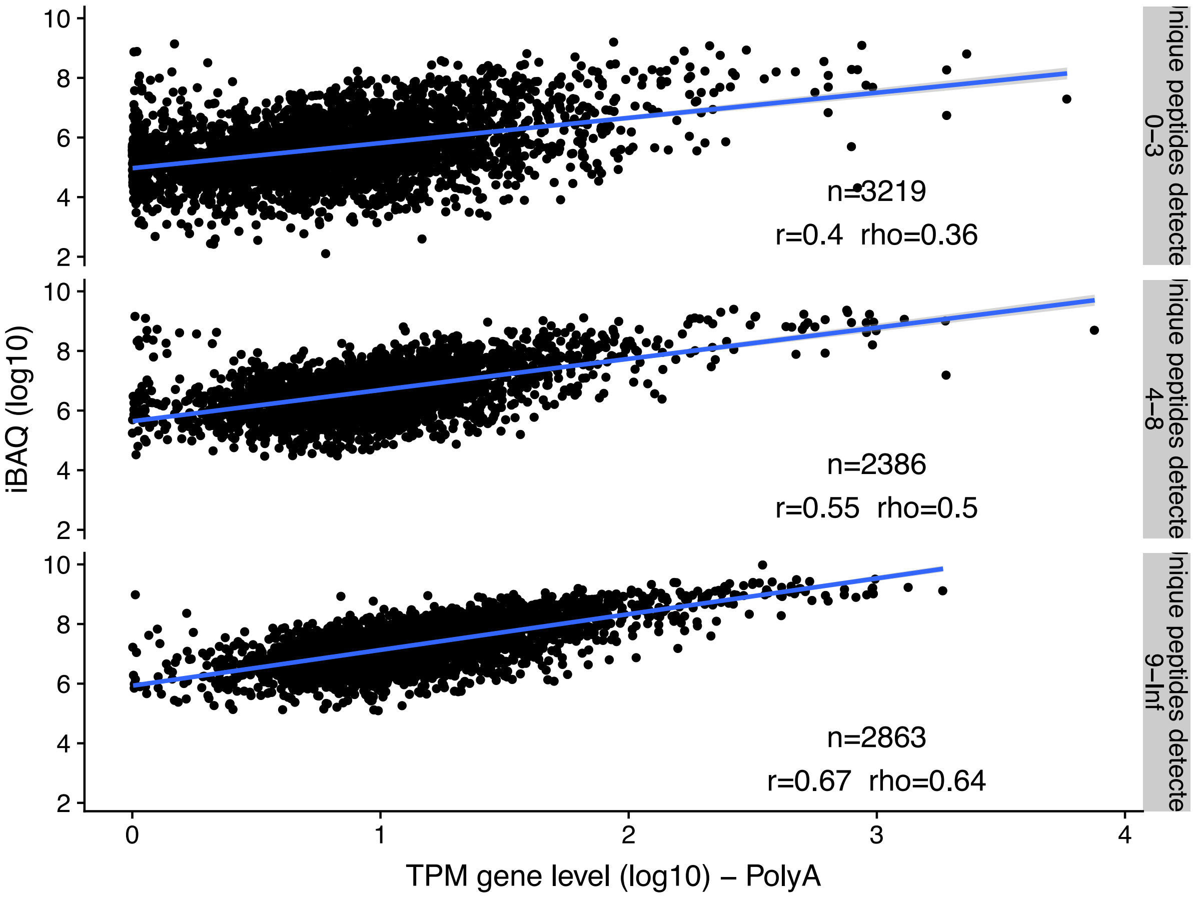


**Supplementary Figure 14:** Agreement between transcript expression and translation with protein abundance. iBAQ values (y-axis) plotted against (x-axis) gene-level TPM values from RNA-seq. Plots are divided according to the number of unique peptides detected. Number of proteins, together with Pearson and Spearman correlations, is shown for each plot.

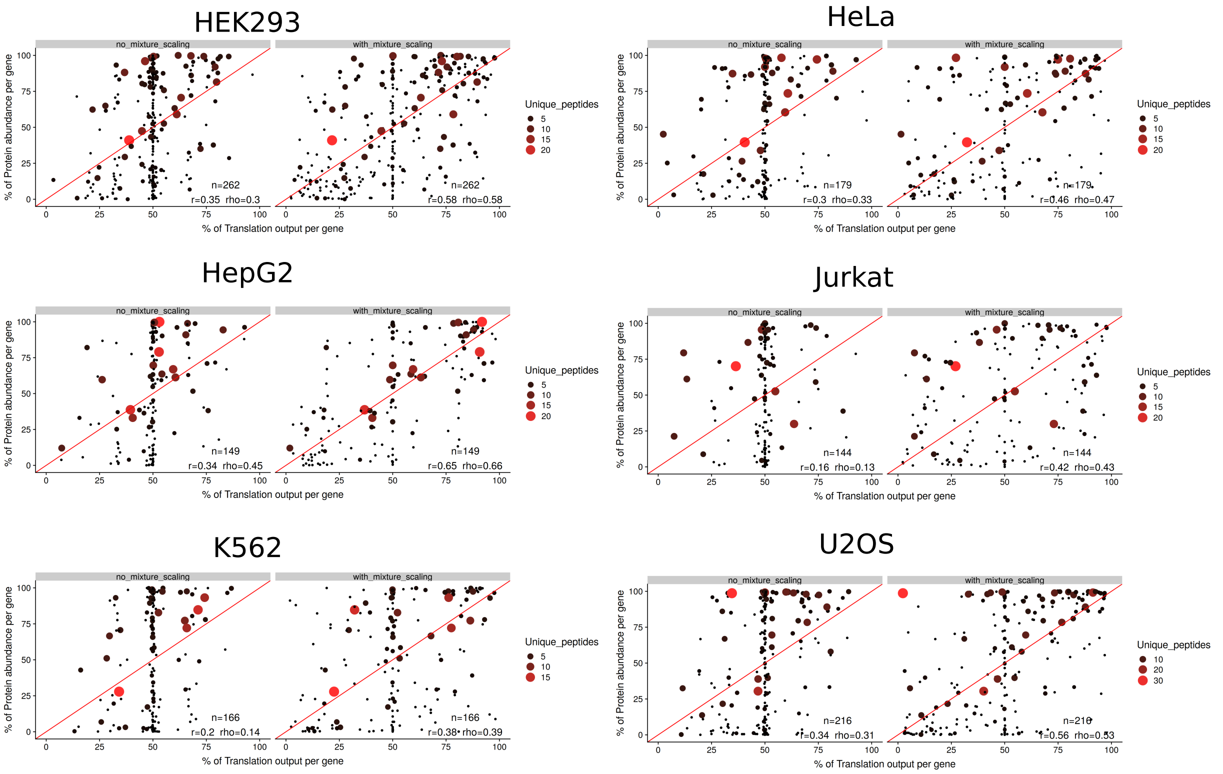


**Supplementary Figure 15:** Agreement between per-gene translation quantification and protein abundance across cell lines, for genes with multiple detected proteins. Pearson and Spearman correlation coefficients between ORFquant-derived % of gene translation and % of gene protein abundance. In each plot, translation quantification is show with (right) and without (left) adjusting for the presence of multiple ORFs.

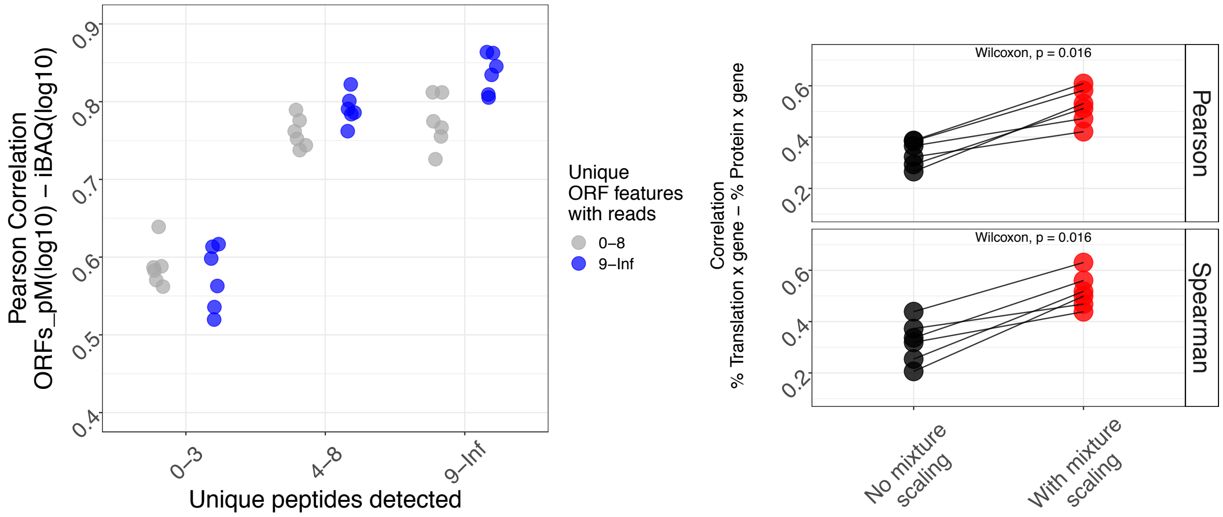


**Supplementary Figure 16:** Correlation between translation quantification and protein abundance using all reads, as shown in Figure 5 using uniquely mapping reads only. Values are shown for all proteins (left), or as percentages of gene signal for protein isoforms (right).
